# Deploying cross-species bone marrow niches to elucidate human hematopoietic stem cell regulation *in vivo*

**DOI:** 10.64898/2026.04.30.719914

**Authors:** Julia Fröbel, Susann Rahmig, Jonas Metz, Iwo Kucinski, Carl-Magnus Svensson, Susanne Reinhardt, Juliane Salbach-Hirsch, Emilie Coppin, Nicole Mende, Nina Henning, Gülce Perçin, Friederike Weschenfelder, Alexandra Köhler, Alexander Platz, Berthold Göttgens, Martina Rauner, Andreas Dahl, Marc Thilo Figge, Thomas Höfer, Claudia Waskow

**Author notes:** Corresponding author Claudia Waskow Immunology of Aging, Leibniz Institute on Aging - Fritz Lipmann Institute (FLI), Beutenbergstrasse 11, 07745 Jena, Germany. Same contribution first authors (Julia Fröbel, Susann Rahmig).

## Abstract

The regulation of human hematopoietic stem cell (HSC) function within its native bone marrow microenvironment remains poorly understood due to the limitations of existing humanized models. Here, we present a non-conditioned human HSC xenotransplantation platform that serves as a physiologically relevant in vivo surrogate to study these complex cellular interactions. Using this system, we uncover a dynamic, cross-species communication between human HSCs and the murine niche, revealing a profound cellular and molecular plasticity of the bone marrow microenvironment in response to humanization. Upon engraftment, platelet-derived growth factor receptor alpha positive (Pdgfra^+^) mesenchymal stromal cells (MSCs) undergo significant numerical expansion and a major transcriptional shift, transitioning from a mixed adipo- and osteo-primed state toward a leptin receptor positive (Lepr^+^) predominantly adipo-primed, hematopoiesis-supporting signature. Functional validation demonstrates that this niche plasticity is a critical determinant of stem cell engraftment: the targeted depletion of Lepr^+^ MSCs or the genetic deletion of stem cell factor results in the rapid mobilization and lack of engraftment of human HSCs, respectively. Our findings establish niche plasticity as a primary regulator of human HSC function and demonstrate the utility of this cross-species platform as a modular genetic toolbox for precise in vivo engineering of the bone marrow microenvironment. Ultimately, this adaptable system provides a powerful framework for elucidating human stem cell biology and advancing bone marrow transplantation therapies.

## INTRODUCTION

Adult hematopoiesis takes place in the bone marrow (BM), where hematopoietic stem cell (HSC) function is coordinated through a specialized, nurturing microenvironment known as the niche. These essential spaces are established around birth (Perçin et al., 2025) and remain vital for lifelong blood homeostasis (Matteini et al., 2021; Schleicher et al., 2024). Our understanding of niche architecture has largely been defined by the discovery of niche cells that balance HSC quiescence and self-renewal against differentiation. Key regulators such as CXCL12 and stem cell factor (SCF), the ligand of the KIT receptor, are provided by a diverse cellular landscape: arteriolar endothelial cells (ECs) and pericytes near the endosteum, alongside *Lepr^+^* mesenchymal stromal cells (MSCs) and sinusoidal ECs within the central marrow cavity, but also CXCL12 abundant reticular cells (CAR) located close to the endosteum, with some of the populations displaying significant overlap (Méndez-Ferrer et al., 2010; Ding et al., 2012; Greenbaum et al., 2013; Mende et al., 2019; Matsuoka et al., 2023). To add to this complexity, mature hematopoietic cells, mainly macrophages of embryonic origin (Jacome-Galarza et al., 2019; Perçin et al., 2025), also play a role in the supportive niches, and orchestrate niche establishment, making them amenable for the seeding with definitive HSCs (Perçin et al., 2025). Ultimately, the stem cell niche is a complex ecosystem whose stability is maintained through a sophisticated interplay between hematopoietic but also non-hematopoietic cells including MSCs and ECs, that both regulate the maintenance or differentiation of stem cells into hematopoietic lineages (Pinho and Frenette, 2019; Fröbel et al., 2021). In this context, the mechanism of human HSC engraftment in mice, including their cellular communication partners within the bone marrow niche that confer niche compatibility between species, is not understood.

Cross-species chimeras, specifically humanized mice, are a powerful model to investigate the function of human HSCs *in vivo*, and have been successfully used to characterize the differentiation propensity of human stem and progenitor cells (HSPCs) (Notta et al., 2011; Cosgun et al., 2014). Moreover, humanized mice have been shown to mount innate and adaptive immune responses from human-derived immune cells (Chupp et al., 2024). However, conventional transplantation regimen rely on toxic conditioning therapy which negatively impacts on the function of human donor cells (Shen et al., 2012; Shao et al., 2014; Hu et al., 2021) and the murine microenvironment (Zhou et al., 2017). To obviate the need for conditioning, we have developed mouse models that robustly engraft human donor HSCs without prior treatment (Cosgun et al., 2014; Mende et al., 2015). As a consequence, the recipient mice support multilineage reconstitution of human blood cells, including myeloid (Cosgun et al., 2014), and erythroid and megakaryocytic lineages (Rahmig et al., 2016; Yurino et al., 2016), but also functional human T cells (Coppin et al., 2021). NSGW41 (NOD.*Prkdc^ko/ko^;Il2rgc^ko/ko^;Kit^W41/W41^*) mice carry a loss-of-function mutation in the KIT receptor mediating a numerically reduced and functionally impaired endogenous murine HSC compartment (Thorén et al., 2008; Waskow et al., 2009), suggesting that human HSCs carrying a fully functional allele of the *Kit* receptor out-compete endogenous, murine HSCs expressing loss of function KIT receptors. We here address this hypothesis experimentally and provide evidence for excellent niche-HSC communication compatibility across the species boundary.

Reciprocal interactions between hematopoietic cells and the niche constitute a fundamental regulatory axis in both, physiological hematopoiesis and leukemogenesis, also contributing to the development of pharmacological resistance (Méndez-Ferrer et al., 2020; Matteini et al., 2021; Schleicher et al., 2024; Urs et al., 2024). Despite the importance of these interactions in human patients, our understanding of the stem cell niche remains largely confined to discoveries from mouse studies, with limited validation in human systems (Chen et al., 2023). To investigate the communication between human hematopoietic stem and progenitor cells and their niche, sophisticated *ex vivo* and *in vivo* platforms have been engineered. These models are reconstituted with human cellular components of the niche to recapitulate the native bone marrow environment (Abarrategi et al., 2018; Dupard et al., 2020). However, so far mainly MSCs and ECs have been integrated in these models, thus reducing niche complexity significantly (Matteini et al., 2021). IPS cell-derived *in vitro* models integrate different hematopoietic and non-hematopoietic cell types of the niche, but they often display incomplete maturation and only limited maintenance of LT-HSCs (Stolz et al., 2026). Further, many cell types display functional heterogeneity *in situ* that is incompletely recapitulated in engineered models. Thus, while current engineered models have laid a great foundation, they inherently simplify the niche, and integration of other hematopoietic or non-hematopoietic niche cell components remains a future challenge. Based on these considerations, mapping the molecular dialogue between the mouse niche and human hematopoiesis provides essential clarity on model translatability and evolutionary conserved mechanisms of niche-HSC cross talk. Finally, these insights act as a catalyst for next-generation experimental designs using humanized mouse models to understand human hematopoietic stem cell biology.

Although *Kit*-mutant mice are indispensable models for studying human HSPCs, the precise mechanisms governing successful human HSC engraftment in the murine environment remain poorly understood. Here, using the NSGW41 strain, we show that long-term human hematopoiesis and secondary reconstitution occur at the expense of endogenous murine stem cells, triggering profound remodeling and transcriptional reprogramming of the MSC microenvironment. To dissect the precise regulators of this phenomenon, we established a highly versatile genetic toolkit, featuring a novel humanized mouse model engineered on a pure C57BL/6 background.

This platform permits the integration of any targeted genetic modification into a murine host supporting human hematopoiesis, offering an unprecedented tool to characterize individual cellular and molecular niche components and their interaction with human HSPCs *in vivo*. Utilizing these novel models, we successfully delineated the specific contributions of the host microenvironment to the regulation of human HSPC function, ultimately identifying LepR⁺ MSC-derived SCF as a primary driver of human HSC engraftment and maintenance *in vivo*.

## RESULTS

### Efficient engraftment of functional human hematopoietic stem cells (HSCs) in NSGW41 mice results in a reduction of endogenous murine HSC numbers

To assess the sensitivity for human HSC engraftment in non-conditioned NSGW41 compared to conventional irradiated NSG mice, titrated numbers of donor human CD34^+^ cord blood hematopoietic stem and progenitor cells (HSPCs) were transplanted. Human leukocyte engraftment in the blood is significantly increased in NSGW41 mice compared to conditioned NSG recipients (**Figure 1A**). Limiting dilution experiments revealed an increased human leukocyte engraftment in the bone marrow (BM) of NSGW41 mice (**Figure 1B**). In NSGW41 the detection threshold is 22-fold higher compared to conditioned NSG recipients as 1/514 (NSGW41) vs. 1/15098 (NSG) donor cells are classified as functional human HSCs based on successful long-term engraftment of human leukocytes (**Figure 1C**). Consistently, frequency and number of human HSCs (Notta et al., 2011; Cosgun et al., 2014) and HSPCs are increased in NSGW41 mice compared to irradiated NSG recipients (**Figures 1D,E, S1A-I**). To evaluate the impact of human donor cell engraftment on endogenous murine stem cells within both recipient strains, HSCs of human and murine origin were quantified (**Figure 1E**). After humanization, the stem cell pool in both recipient strains increases in absolute numbers, however, the composition differs strikingly. In irradiated NSG, HSCs of human and murine origin are present in equal numbers, whereas human HSCs outnumber murine HSCs in NSGW41 recipient mice by far, suggesting a distinct mechanism of engraftment. Endogenous murine HSCs are significantly reduced in NSGW41 but not in irradiated NSG mice. The effect remains significant at the level of hematopoietic progenitors (**Figure S1J**), suggesting that in NSGW41 mice, Kit-deficient, endogenous HSPCs are outcompeted by human KIT-proficient donor cells. Overall, increased engraftment efficiency in NSGW41 mice compared to irradiated NSG mice is consistent with the elevated numbers of functional, growth-factor responsive human progenitors harboring the propensity to differentiate into myeloid, erythroid and megakaryocytic lineages in NSGW41 (Rahmig et al., 2016). To quantify engraftment of functional human HSCs in primary recipients, titrated numbers of human CD45^+^ BM cells isolated from primary recipients were transplanted into secondary recipient mice (scheme, **Figure 1F**). Secondary transfer of human BM leukocytes from NSGW41 mice resulted in substantially greater engraftment than those sourced from irradiated NSG primary recipients. (**Figure 1G**). Moreover, while mean leukocyte chimerism in the bone marrow of secondary irradiated NSG recipients improves only marginally with increased donor cell numbers, secondary NSGW41 recipient mice prove highly amenable to engraftment of titrated donor cells from primary recipients, demonstrating dose-dependent engraftment levels. The frequency of human HSCs successfully repopulating secondary recipient mice is increased ∼5-fold in primary NSGW41 compared to primary irradiated NSG mice (**Figure 1H**). Phenotypic as well as functional identification of human donor HSCs in primary recipients shows an advantage of NSGW41 over irradiated NSG recipients. Specifically, the total number of functional human HSCs engrafted in primary recipients increases by 14-fold using NSGW41 compared to irradiated NSG mice, as assessed by secondary transplantation capacity (**Figure 1I**). This highlights the extent of stable human stem cell engraftment in NSGW41 mice.

**Figure 1:**
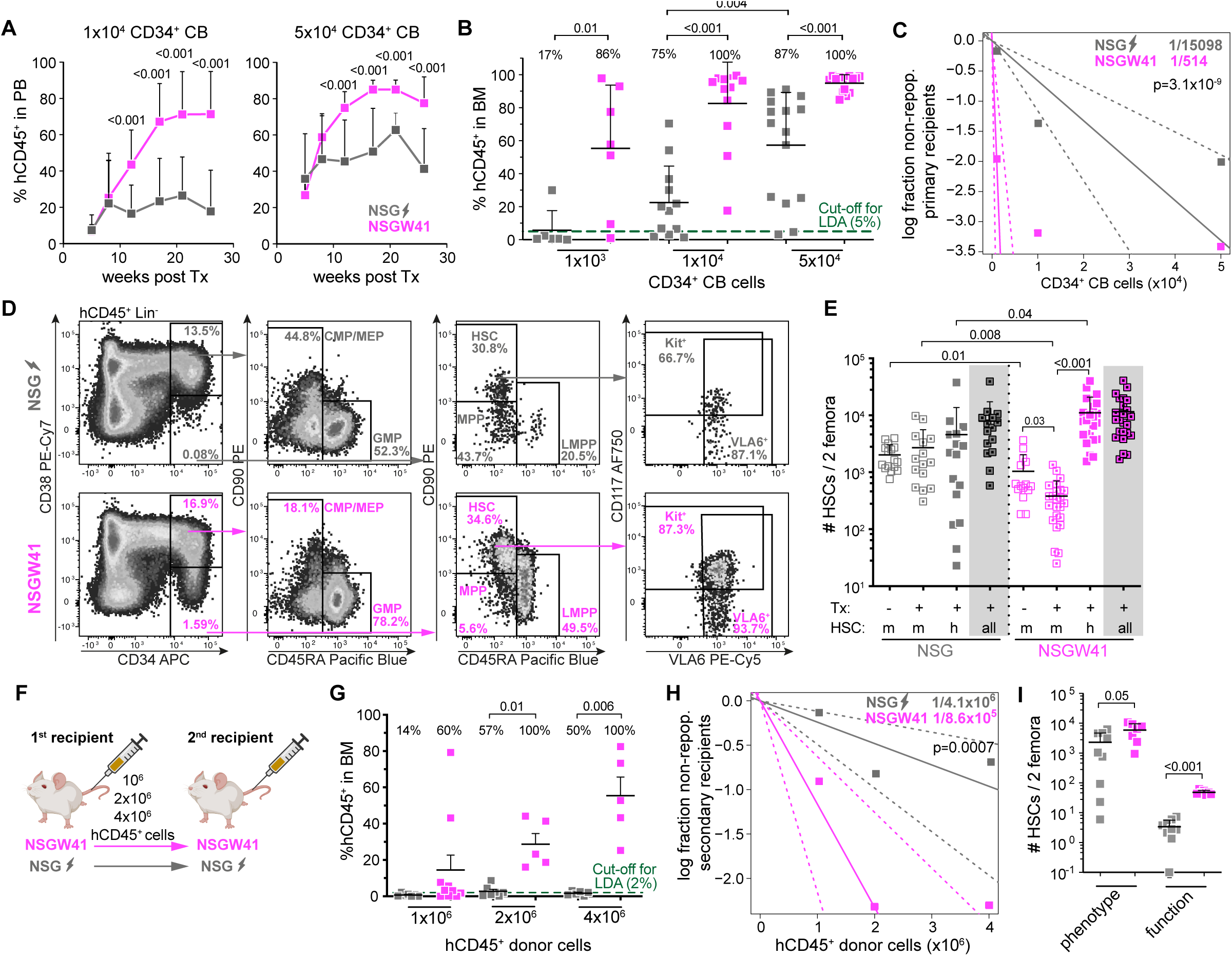
Near complete engraftment and maintenance of functional human HSCs in NSGW41 mice. **A** Repopulation kinetic in the blood of non-irradiated NSGW41 (pink) and irradiated NSG (grey) recipient mice that have received 1×10^4^ (left) or 5×10^4^ (right) enriched CD34^+^ cord blood (CB) cells. Data summarized from 4 or 3 independent experiments using 13 (1×10^4^ CB cells) and 16 or 21 (5×10^4^ CB cells) recipient mice, respectively. PB-peripheral blood. **B** Indicated titrated numbers of CB cells were transplanted into non-irradiated NSGW41 and irradiated NSG mice. Contribution of human CD45^+^ cells to BM leukocytes was analyzed 24-35 weeks later. Percentages on top indicate the frequency of mice that showed successful engraftment (>5% of human leukocytes in the BM). BM-bone marrow, LDA-limiting dilution analysis. **C** Extreme limiting dilution assay (ELDA) calculations based on data shown in B. Human leukocyte engraftment in the bone marrow above 5% at 24-35 weeks after transplantation was evaluated as positive engraftment. Stem cell frequencies are significantly different between both recipient lines, p=3.1×10^-9^. **D** Representative FACS plots of human donor-derived HSPCs in irradiated NSG (top) and non-irradiated NSGW41 mice (bottom) 26 weeks after humanization. **E** HSC numbers of murine (KSL CD48^-^CD150^+^, open symbols) and human (CD34^+^CD38^-^CD90^+^CD45RA^-^, closed symbols) origin, or added together (grey background) in mice as indicated with and without prior humanization. Analysis was performed 24-29 weeks after transplantation and in age-matched non-transplanted controls. Statistics between different mice were calculated by Welch’s t-test, for different cells within the same mice by paired t-tests. **F** Scheme for secondary transplantation. Indicated titrated numbers of sorter-purified human CD45^+^ bone marrow cells from primary recipient mice (irradiated NSG or non-irradiated NSGW41) were transplanted into secondary recipients of the same strain and treatment and human leukocyte chimerism in the BM was analyzed 24-27 weeks post transplantation. **G** Human leukocyte chimerism in the bone marrow of secondary recipients. Percentages on top indicate the frequency of mice that showed successful engraftment (>2% of human leukocytes in the BM). **H** ELDA calculations based on data shown in G. Engraftment above 2% was defined as positive engraftment. Stem cell frequencies are significantly different, p=7×10^-4^. **I** Numbers of human HSCs in primary recipient mice determined by phenotype (flow data) versus functional assessment (ELDA in secondary recipients).

### Human HSC pool size correlates with increased numbers of mesenchymal stromal cells

To test for a response of the stem cell niche to humanization, we determined the number of non-hematopoietic niche cells in the BM and endosteum (Mende et al., 2019; Perçin et al., 2025) (**Figures 2A-D**). BM-resident, but not endosteal mesenchymal stromal cells (MSC, CD51^+^Pdgfra^+^Sca1^-^) significantly increase in NSGW41 mice in response to humanization (**Figure 2C**). MSC expansion is only detected in transplanted, non-irradiated NSGW41 mice but not in irradiated NSG recipients, suggesting a specific role for this cell type in efficient repopulation with human HSCs in NSGW41 mice. In contrast, osteoblasts or endothelial cells are numerically not altered compared to control non-transplanted NSGW41 mice. BM resident endothelial cells are reduced in humanized NSG mice, likely based on incomplete recovery after the irradiation insult. Transplanted NSGW41 mice exhibited sustained MSC expansion, maintaining significantly higher niche compartment counts than age-matched controls throughout a ∼30-week observation period (**Figure 2E**). In humanized NSGW41 mice, donor HSC numbers exhibit a significant positive correlation with endogenous MSC numbers, but neither with endothelial cells nor osteoblasts in the BM (**Figure 2F**). The positive correlation is only seen in BM but not the endosteal bone region (**Figure S2**). Together, this data suggests that the cellular mouse niche responds to humanization with a specific expansion of MSCs, possibly amplifying available niche space or factors to host human and murine HSCs. We conclude that the niche response to humanization differs between NSGW41 and irradiated NSG mice and that MSCs specifically expand in NSGW41 mice. This expansion may be a prerequisite for efficient long-term engraftment of human donor HSCs observed in NSGW41.

**Figure 2:**
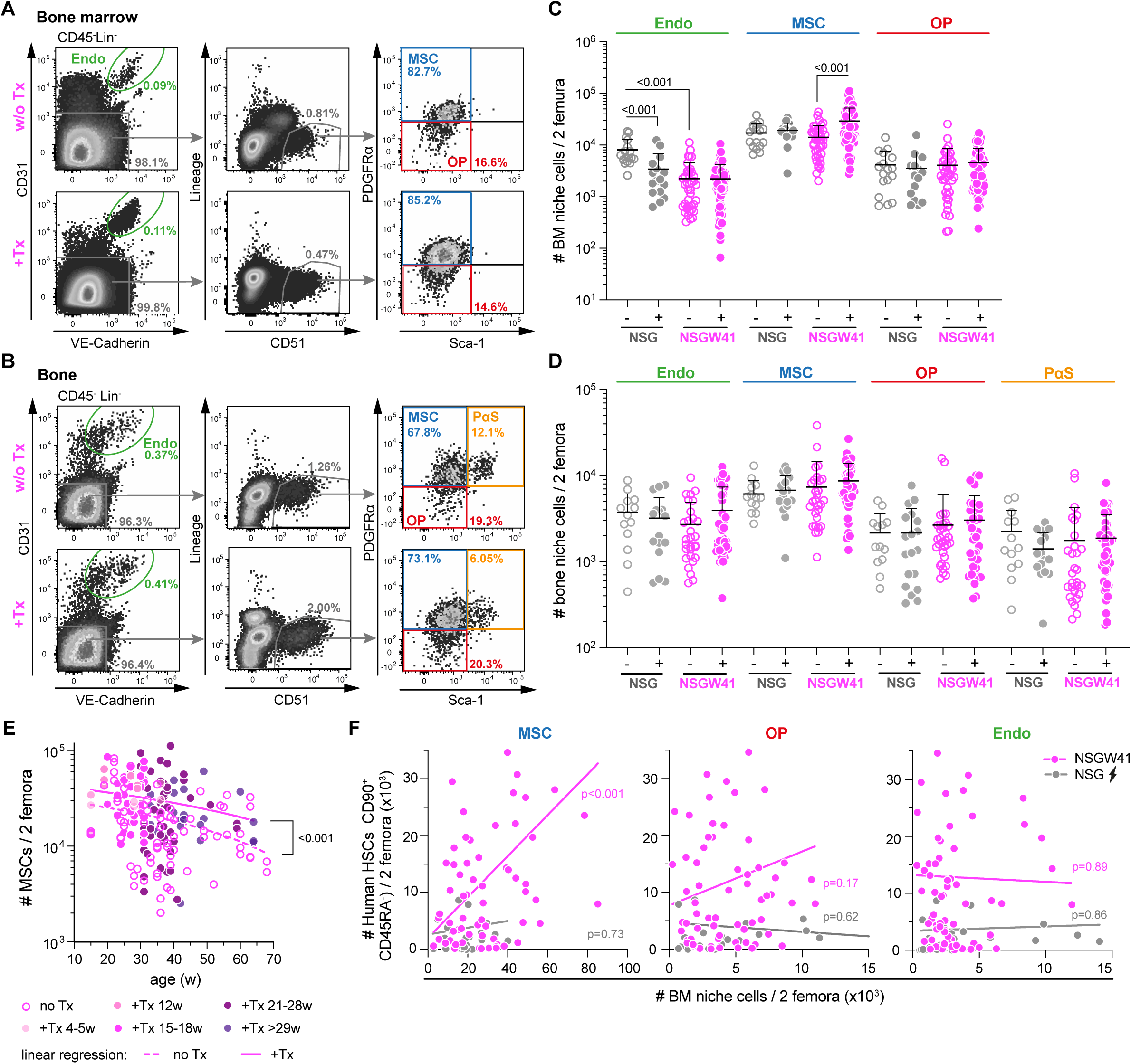
Cellular response of bone marrow niche to the transplantation of human HSCs. A,B. Representative FACS profiles showing the identification of endothelial cells (Endo), CD51^+^PDGFRɑ^+^ mesenchymal stromal cells (MSC), and CD51^+^PDGFRɑ^-^ osteoprogenitors (OP) in the bone marrow (A) and the same cells plus CD51^+^PDGFRɑ^+^Sca-1^+^ PaS cells in the bone (B) fractions of untreated control mice and humanized mice that have received human donor CB cells 29 weeks before. **C,D** Quantification of niche cell populations as shown in A and B in the bone marrow (C) and bone (D) fractions of 2 femora. Data of 14 independent experiments analyzed 23-29 weeks post transplantation is shown. **E** MSCs per 2 femora from non-humanized and humanized NSGW41 mice plotted against age. Data were analyzed by linear regression. Intercepts of humanized NSGW41 MSC numbers were found to be significantly elevated throughout the test period of >50 weeks after confirming that the slopes were statistically indistinguishable (color code indicates weeks post Tx). **F** Correlation between the numbers of bone marrow murine MSCs (left), osteoprogenitors (middle), or endothelial cells (right) and engrafted human HSCs (15 independent experiments, 4-35 weeks post Tx, p-values of Pearson correlation).

### Plastic molecular response of niche MSCs towards adipogenic potential

To elucidate whether, in addition to quantitative changes, niche MSCs also undergo molecular adaptations in response to humanization, a comparative transcriptome sequencing analysis was conducted. To this end, sorter-purified CD51^+^Pdgfra^+^Sca1^-^ MSCs were isolated from humanized NSG (irradiated) and NSGW41 mice, alongside their respective non-humanized controls. Principal component analysis revealed distinct transcriptional profiles in MSCs isolated from NSG or NSGW41 mice before and after humanization (**Figure 3A**). Specifically, humanized MSCs from both strains cluster together in the PC-space. This suggests a general independent – transcriptional adaptation of MSCs in response to humanization which is further indicated by the large number of genes (>1500) that are consistently deregulated by humanization in both genotypes (**Figure 3B**, middle bar, **Table S3**). To link the observed transcriptional changes to molecular processes in MSCs that were modulated by humanization, we conducted an unbiased gene set enrichment analysis (GSEA). We detect an adaptive immune signature in NSG mice that is absent in MSCs from humanized NSGW41 mice (lymphocyte activation, **Figures 3C, S3A,B, Table S4**). Except for the NSG-specific immune response, all gene sets significantly affected by humanization in at least one mouse line mainly show reduced gene expression, with similar changes observed in the other mouse strain as well (**Figures 3C, S3A,B, Table S4**). These deregulated gene sets reflect coordinated reductions in the expression of genes involved in skeletal system development, ossification, and bone or cartilage formation (**Figures 3C, S3A,B**, **Table S4**). This finding highlights a molecular response of MSCs to incoming human donor HSCs, possibly affecting MSC cell fate and therefore also the capability of MSCs to support human hematopoiesis.

**Figure 3:**
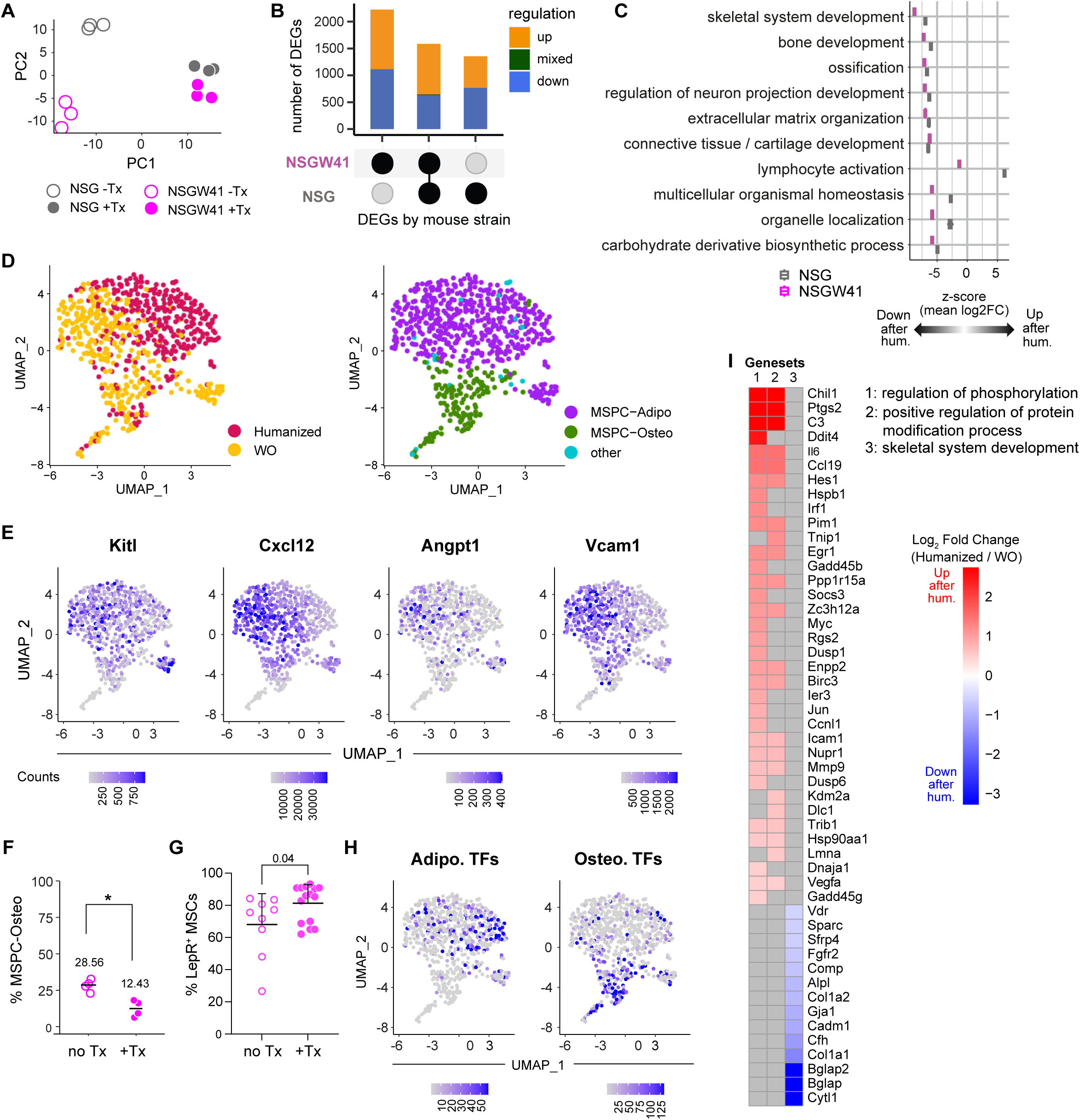
Reduced osteogenic transcriptional priming in MSCs upon humanization. **A** PCA plot of bulk RNA sequencing data of sorter-purified MSCs from irradiated NSG or NSGW41 with (Tx) or without (-Tx) humanization. 500 most variable genes were used. **B** Mouse line-specific quantification of genes differentially expressed in response to humanization. **C** Summary of Gene Set Enrichment Analysis (GSEA) showing magnitude and direction of pathway-level expression changes in MSCs from NSG and NSGW41 mice after humanization. Each row represents a functional module of related GO Biological Process terms (“umbrella term”). For each genotype, average gene expression changes (z-scores of log₂ fold changes) across the GO terms in that module are shown, providing a sense of the overall strength and direction of pathway regulation. For expression changes of individual genes within these modules see **Figure S3B**. **D** UMAP plots of single-cell transcriptomes of sorter-purified MSCs from humanized and non-humanized NSGW41 mice(both n=4). MSCs were extracted from humanized mice 24 weeks after transplantation. Cells are colored by condition (left) and MSC-subtype (right). For subtype annotation see **Figure S3C,D**. **E** Normalized transcript counts of selected genes known to support hematopoiesis. **F** Fraction of osteo-primed single-cell transcriptomes of all MSC transcriptomes per mouse (p-value <0.05, Wilcoxon test). **G** Frequency of *LepR*-expressing MSCs as determined by flow cytometry from humanized NSGW41 mice (15-28 weeks) and non-humanized controls (student’s t-test). **H** Seurat expression module scores of lineage associated transcription factors: Adipogenesis: *Cebpb*, *Cebpd*, *Pparg*, *Cebpa*; Osteogenesis: *Runx2*, *Sp7*. For expression patterns of individual TFs see **Figure S3F**. **I** Humanization-induced expression changes in gene modules identified by GSEA of single-cell transcriptomes. Gray fields indicate that genes are not included in the functional module (“umbrella term”) indicated by the column name. GO terms summarized by these umbrella terms are shown in **Figure S3G**.

To assess whether all MSCs uniformly alter their transcriptional profile after humanization or whether a MSC subpopulation expands and finally dominates the transcriptional data, we performed single-cell sequencing of MSCs from humanized NSGW41 and control mice. In a two-dimensional UMAP embedding of single-cell transcriptome profiles, clusters appear distinctively populated by cells from humanized and non-humanized mice but experimental batches have no apparent impact on the distribution of cells (**Figures 3D left, S3C**). This indicates technical neutrality and suggests that humanization evoked transcriptional alterations (**Figures 3D, left**). To evaluate if the distinct distribution of MSCs from humanized mice in the UMAP embedding reflects a shift in the prevalence of MSC subpopulations, we projected the RNA-profiles of our cells onto previously published niche single cell data sets (summarized in ((Dolgalev and Tikhonova, 2021) (**Figure S3D**). Based on this projection we could confirm the MSC identity of our samples and could further subclassify our MSCs into adipogenic mesenchymal stromal progenitor cells (MSPC-adipo) and osteogenic mesenchymal stromal progenitor cells (MSPC-osteo) that form clearly separated clusters in the UMAP embedding (**Figure 3D, right**). Specifically, those of our MSC profiles with high expression of *Lepr*, *Cxcl12*, *Adipoq* and *Lpl* are annotated as MSPC-adipo whereas those with high expression of *Bglap* and *Spp1* are annotated as MSPC-osteo (**Figure 3D right, S3E**). In contrast to osteogenic MSPCs, adipogenic MSPCs also express higher levels of the stem cell supporting factors *Kitl*, *Cxcl12*, *Angpt1*, and *Vcam1* (**Figure 3E**). The single-cell MSC profiles show two adaptations upon humanization: While the fraction of MSCs annotated as osteogenic decreases (**Figure 3D,F**), conversely meaning that the fraction of adipogenic MSPCs increases, the MSPC-adipo from humanized mice show reduced transcript levels of *Cxcl12*, *Angpt1*, and *Vcam1* (**Figure 3E, Table S5**). The increase in MSPC-adipo is consistent with flow cytometry results which confirm an increase in Lepr-positive MSCs in BM niches of humanized NSGW41 mice (**Figure 3G**). Next, we asked if transcriptional changes in MSPC-adipo upon humanization indicate changes in their lineage-propensity. Therefore, we scored cells by transcript levels of adipo-and osteogenic transcription factors (TFs) (**Figure 3H, S3F, Table S5**). Transcript levels of adipogenic TFs, especially *Cebpa* and *Cebpb* (**Figure S3F**), are increased in the region in the UMAP that is predominantly populated by MSPC-adipo from humanized mice. In contrast, transcripts of osteogenic TFs are increased in the region in the UMAP where MSPC-osteo from control and humanized cells are located (**Figure 3H, S3F**). Together, this suggests that upon humanization an increased fraction of MSCs acquires the transcriptional state of stem cell-supporting adipogenic MSCs, while the fraction of MSCs with osteogenic potential is reduced. This shift in lineage-propensity appears to be the dominant effect of humanization that is also reflected by an unbiased GSEA. Consistent with the analysis of bulk RNA-seq data the analysis reveals a reduction of MSC-transcripts mapped to the GO pathway skeletal system development (**Figure 3I, S3G, Table S6**). We conclude that MSCs after humanization have reduced osteogenic potential but express key factors for supporting the HSC function suggesting that *Lepr^+^* adipogenic MSCs may provide the key niche cell for the engraftment and maintenance of xenogenic human HSCs in mice.

### Physiological relevance of transcriptome alterations

We next investigated whether alterations in the transcriptional profile of MSCs translate into modified differentiation potential of MSCs isolated from humanized or non-humanized mice. The frequency of stromal colonies (CFU-F) generated *in vitro* from sorter-purified MSCs is comparable between humanized NSGW41 or control mice (**Figure 4A**). Also, individual colonies are of comparable sizes, suggesting that MSC expansion *in situ* occurs without major interference with their per-cell propensity to form stromal colonies. Notably, humanization also has no influence on the overall adipogenic differentiation potential of bone marrow cells from NSGW41 mice, irrespective of their humanization (**Figure 4B**). In contrast, and consistent with transcriptional alterations (**Figure 3**), osteogenic differentiation potential decreases after humanization compared to non-humanized controls (**Figure 4C,D**). While general bone morphometric values remain unaltered between humanized and control mice (**Figure 4E**), in-depth analysis reveals a reduced bone mineral density of long bones after humanization (**Figure 4F**). A linear correlation is observed between the decrease in bone mineral density and the increase in MSC numbers, further reinforcing the inverse relationship between these two parameters (**Figure 4G**). This suggests that transplantation induces a physiologically relevant shift in the differentiation potential of MSCs *in vivo*. In patients undergoing hematopoietic stem cell transplantation, bone loss frequently occurs (Schulte and Beelen, 2004), positively validating NSGW41 mice as an excellent and translationally relevant xenotransplantation model.

**Figure 4:**
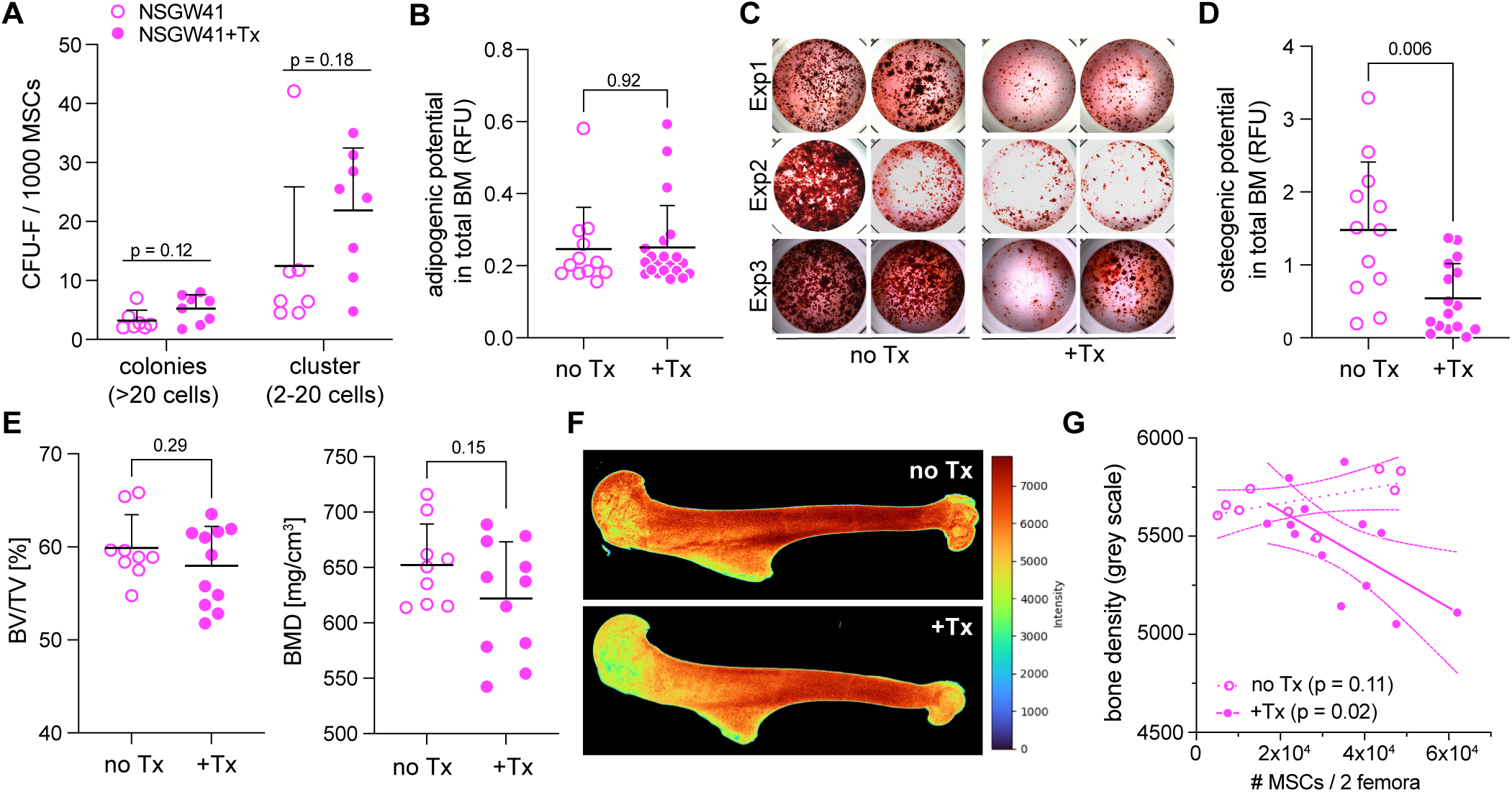
Human HSC engraftment results reduced osteogenic differentiation potential of murine MSCs. **A** Quantification of CFU-F potential of sorter-purified MSCs from humanized NSGW41 mice (4 independent experiments, 15-27 wks post Tx) and non-transplanted controls. CFU-F formation was assessed 16 to 20 days after seeding. Open symbols represent non-humanized controls and closed symbols represent humanized mice. Each dot is derived from one mouse. P-values were calculated by multiple Mann-Whitney tests. **B** Quantification of OilRed O stained adipogenic differentiation of NSGW41 bone marrow cells with and without prior humanization (absorbance at 510 nm). One dot represents cultures from one mouse, 4 independent experiments, 18-26 wks post Tx. **C** Alizarin Red stained osteogenic potential of bone marrow cells from NSGW41 mice with and without prior humanization (26-32 wks post Tx) cultured in osteogenic differentiation medium for 14 days. Representative images from six mice per group are shown. **D** Quantification of Alizarin Red stained calcium deposits as shown in C (absorbance at 450 nm, 4 independent experiments, 26-32 wks post Tx). **E** Bone volume over total volume (BV/TV) and bone mineral density (BMD) from microCT scans of humeri from NSGW41 with and without prior humanization (4 independent experiments; 12-35 wks post Tx). **F** Rainbow-color mapped microCT scans showing bone density of humeri from NSGW41 with and without prior humanization (12 wks post Tx). **G** Mean bone density (measured in humeri) plotted versus the number of MSCs (per 2 femora) in NSGW41 mice with and without prior humanization (12-35 wks post Tx, p-values of Pearson correlation).

### Human HSCs are maintained by *Lepr^+^* MSCs in the bone marrow of humanized NSGW41 mice

To investigate the mechanisms by which *Lepr^+^* MSCs support the engraftment of human HSCs, we engineered a novel humanized mouse model on a C57BL/6 genetic background. This approach allows for the seamless integration of established murine genetic tools to interrogate niche-human HSC interactions *in situ*. By combining the *Rag2^-/-^*, *Il2rg^-/-^*, and *Cd47^-/-^* genotypes with the *Kit^W41/W41^* mutation, we generated *Rag2^-/-^;Il2rg^-/-^;Kit^W41/W41^;Cd47^-/-^*(B6.RgW41CD47) mice. Following the transplantation of ∼5×10^4^ human CD34^+^-enriched cord blood cells, B6.RgW41CD47 mice demonstrate robust humanization. Chimerism levels in the peripheral blood (**Figure 5A**) and bone marrow (**Figure 5B**) are comparable to those observed in NSGW41 mice and significantly superior to irradiated NSG recipients. Furthermore, B6.RgW41CD47 mice exhibit an increased frequency of short-lived human myeloid cells in the BM, a phenotype mirroring that of NSGW41 mice and indicating stable, multi-lineage donor engraftment (**Figure 5C**). Consistent with these observations, the absolute numbers of human HSCs in the BM of B6.RgW41CD47 and NSGW41 recipients are comparable, and both strains show marked increases over irradiated NSG controls (**Figure 5D**). Crucially, B6.RgW41CD47 mice recapitulate the expansion of the MSC compartment upon humanization (**Figure 5E**). These findings validate the reproducibility of HSC-induced MSC expansion across different genetic backgrounds and establish this strain as an ideal platform for functional *in vivo* studies of the humanized hematopoietic niche.

**Figure 5:**
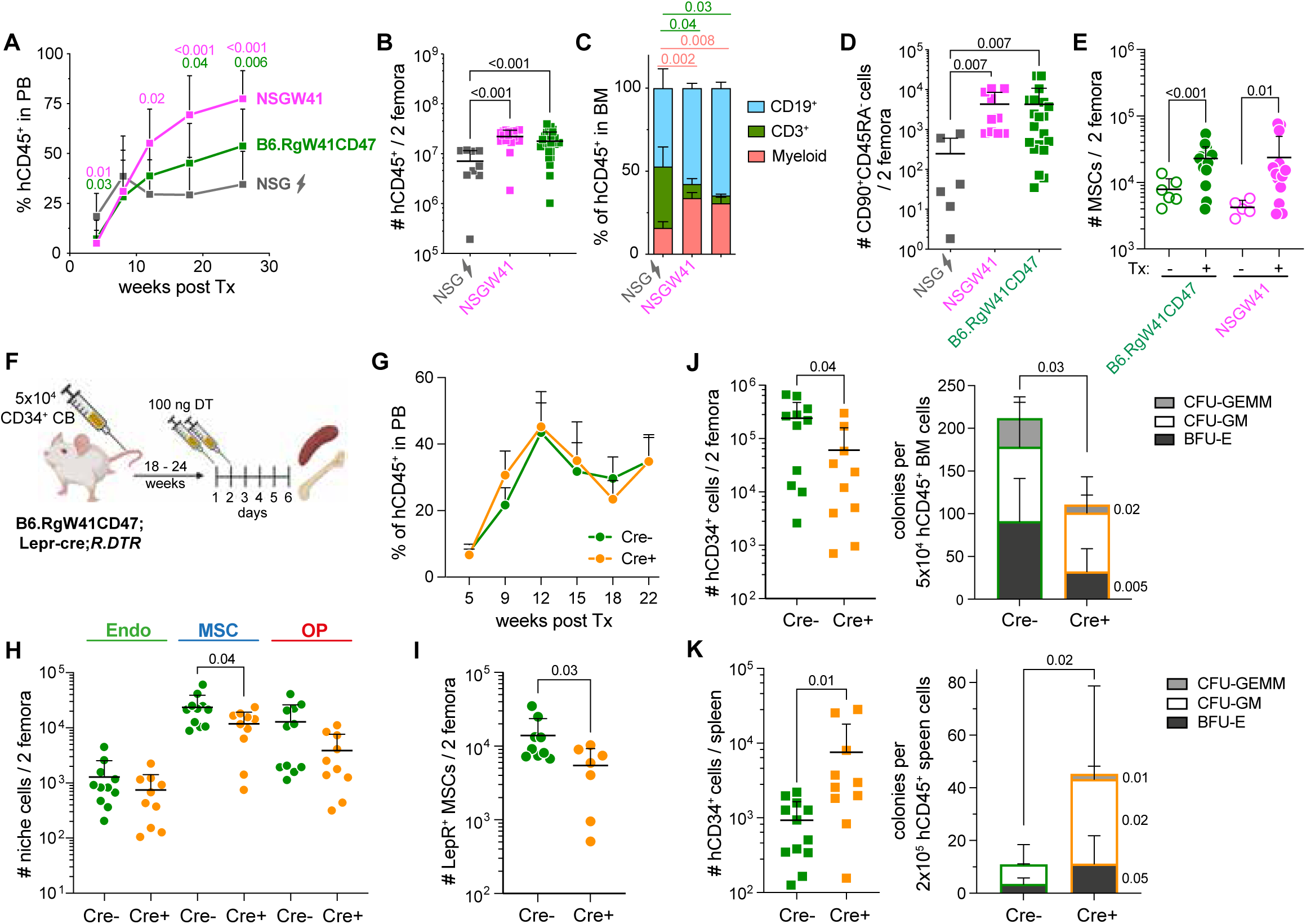
Depletion of *Lepr^+^* MSCs from humanized mice mobilizes human HSPCs to the spleen. **A** Repopulation kinetic of non-irradiated B6.RgW41CD47 and NSGW41 and conditioned NSG recipient mice that have received 5×10^4^ CB cells. Data summarized from 7 independent transplantations into a total of 33 RgW41CD47, 32 NSGW41 and 10 irradiated NSG recipient mice. **B** Number of human CD45^+^ leukocytes in 2 femora of indicated recipient mice 26 to 40 weeks after transplantation. **C** Relative contribution of human HSC-derived T cells (CD3), B cells (CD19), and myeloid cells (CD33 and CD16) in the same mice as in B. **D** Number of human CD34^+^CD38^-^CD90^+^CD45RA^-^ HSCs per 2 femora in indicated mice as outlined in B. **E** Number of CD51^+^PDGFRɑ^+^ MSCs in 2 femora of non-transplanted (-Tx) and transplanted (+Tx) RgW41CD47 and NSGW41 mice. **F** Experimental outline of Lepr^+^ MSC depletion in B6.RgW41CD47;Lepr-cre;R.DTR mice by diphtheria toxin (DT). **G** Repopulation kinetic of non-irradiated B6.RgW41CD47;Lepr-cre;R.DTR recipient mice (11 Cre^-^ and 10 Cre^+^) that have received CB cells from 4 independent donors. **H** Number of endothelial cells, MSCs and osteoprogenitor cells 4 days after the last DT injection in the same mice as in G. **I** Number of *Lepr^+^* MSCs 4 days after the last DT injection in the same mice as in H. **J, K** Left: Quantification of human HSPCs in BM (J) and spleen (K) of Cre^-^ and Cre^+^ B6.RgW41CD47;Lepr-cre;R.DTR mice 4 days after last DT injection. Right: Colony forming potential per 5×10^4^ hCD45^+^ cells from bone marrow (J) or 2×10^5^ seeded hCD45^+^ cells from spleen (K) of Cre^-^ and Cre^+^ B6.RgW41CD47;Lepr-cre;R.DTR mice 4 days after last DT injection.

To evaluate the functional requirement of *Lepr^+^* MSCs in maintaining human HSCs, we generated a conditional depletion model on the B6.RgW41CD47 background (B6.RgW41CD47;Lepr-cre;iDTR; **Figure 5F**). Engraftment kinetics between B6.RgW41CD47;Lepr-cre;iDTR and control mice are identical (**Figure 5G**) and diphtheria toxin (DT) administration results in the targeted ablation of MSCs without affecting other niche populations (**Figure 5H**). Moreover, a specific depletion of *Lepr^+^* cells within MSCs is detected (**Figure 5I**). The depletion of *Lepr^+^* MSCs leads to a significant reduction in BM-resident human CD34^+^ donor cells (**Figure 5J, left**). The loss of CD34^+^ HSPCs is accompanied by a diminished colony-forming unit (CFU) capacity among sorter-purified human bone marrow leukocytes, suggesting that both primitive stem cells and downstream progenitors are lost or mobilized from the marrow in the absence of *Lepr^+^* MSCs (**Figure 5J, right**). In support of the mobilization hypothesis, we observed a concomitant increase of human CD34+ HSPCs and growth factor-responsive progenitor numbers in the spleens of *Lepr^+^* MSC-depleted mice (**Figure 5K**). Specifically, while CFU-GEMM and BFU-E frequencies, but not CFU-GM, are reduced in the bone marrow, all colony types are significantly enriched in the spleen, indicating a preferential migration of multipotent and erythroid-biased progenitors to the splenic environment. Collectively, these data demonstrate that the depletion of murine *Lepr^+^* MSCs triggers a shift of human HSPCs from the BM to the periphery, providing direct evidence that *Lepr^+^* MSCs are essential for the retention and maintenance of human HSCs within the xenogeneic BM niche.

### *Lepr^+^* MSC-derived murine SCF is crucial for the tethering of human donor HSCs in the stem cell niche

SCF is a critical regulator of murine HSC biology and *Lepr^+^* MSCs are the major source of stem cell factor (SCF) in the BM (Waskow et al., 2002; Ding et al., 2012). Despite a reported 100-fold reduction in binding affinity for the human KIT receptor (Lev et al., 1992), murine SCF efficiently supports the growth of human CD34^+^ cord blood-derived HSPCs *in vitro* (**Figure 6A**). This finding may be explained by high protein similarity of ligands and receptors between humans and mice (>80%, (Fröbel et al., 2021)), indicating functional conservation of the SCF-KIT signaling axis across species. We hypothesize that human HSCs replace mouse HSCs based on their competitive binding and consumption of murine MSC-derived SCF. Consistent with this idea, while hypomorphic levels of KIT are expressed by HSCs from non-transplanted NSGW41 mice, after humanization only murine HSCs which express normal levels of KIT-receptor expression remain (**Figure 6B**). In HSCs from NSG mice we see no difference in the level of KIT expression with and without humanization. The bioavailability of soluble SCF significantly decreases in the bone marrow but not in the blood of humanized NSGW41 mice (**Figure 6C**). These findings further support the hypothesis that the Kit-SCF axis is involved in the engraftment process of human HSCs in mice.

**Figure 6:**
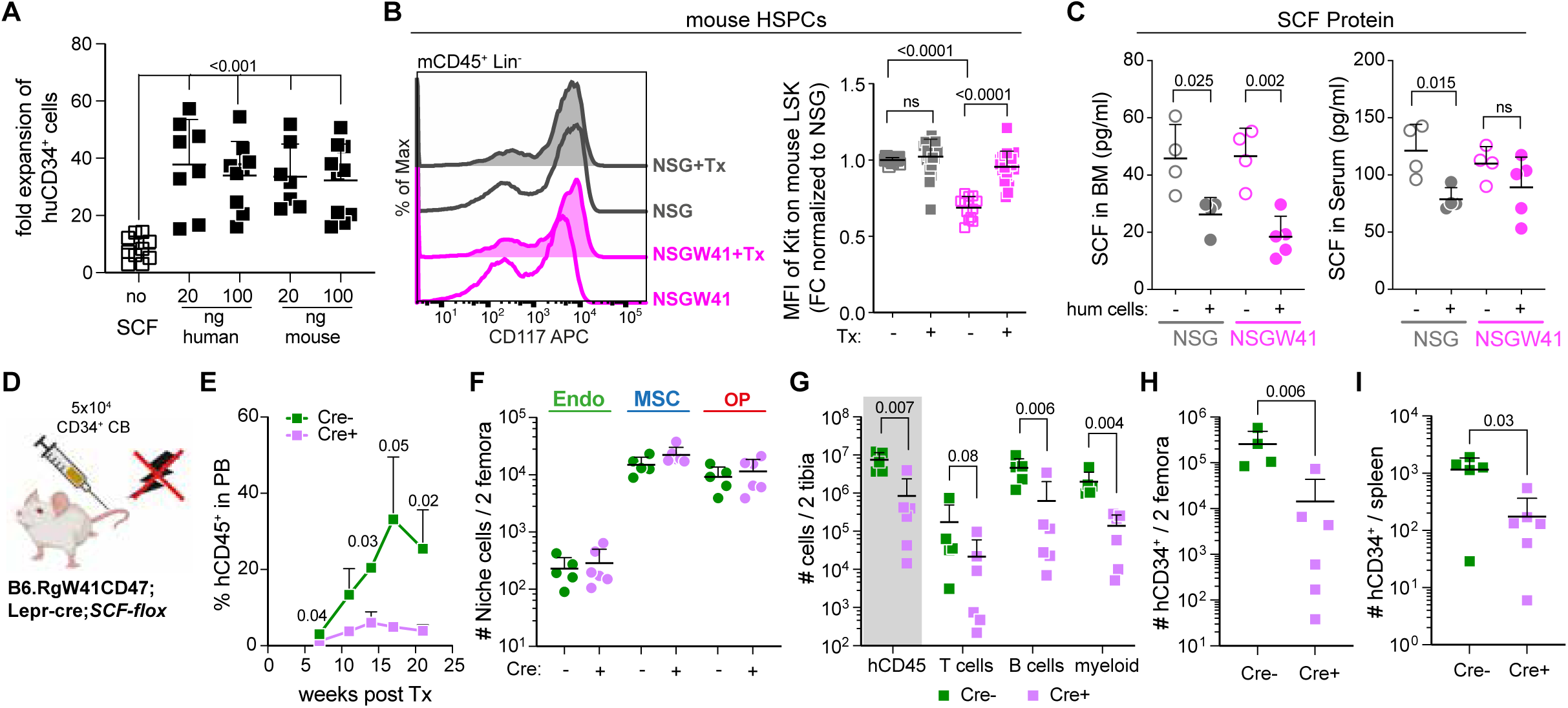
MSC-derived SCF is crucial for human HSC engraftment. **A** Response of human CD34^+^ cord blood cells to human or mouse stem cell factor (SCF) *in vitro*. Depicted is the fold-expansion after 7 days of culture (6 independent experiments, ordinary one-way ANOVA). **B** Representative histograms showing KIT expression on mCD45^+^Lin^-^ BM cells from a humanized and a non-humanized NSG and NSGW41 mouse (left). Fold change of mean fluorescence intensity (MFI) of KIT on mCD45^+^Lin^-^ cells from humanized (24-35 weeks) and non-humanized NSG and NSGW41 mice normalized to non-humanized NSG mice, 3 independent experiments (right). P-values were calculated by unpaired t-test. **C** Plots show SCF concentrations in the bone marrow (left) and serum (right) of NSGW41 and NSG mice with and without humanization. P-values were calculated by student’s t-tests. **D** Scheme of experimental approach for humanization of B6.RgW41CD47;Lepr-cre;SCF-flox recipient mice. **E** Repopulation kinetic of non-irradiated B6.RgW41CD47;Lepr-cre;SCF-flox recipient mice and controls (5 Cre^-^ and 6 Cre^+^) that have received CB cells. Two independent experiments. P-values were calculated by multiple unpaired t-tests. **F** Number of endothelial cells (Endo), mesenchymal stromal cells (MSCs) and osteoprogenitor cells (OP) 21 weeks after humanization of the same mice as in F. **G** Number of mature human leukocytes 21 weeks after humanization in the bone marrow of the same mice as in F. **H, I** Number of human CD34^+^ HSPCs 21 weeks after humanization in the bone marrow (I) and spleen (J) of the same mice as in F-H.

To test for functional significance of MSC-derived murine SCF for the engraftment of human donor cells, B6.RgW41CD47;Lepr-cre;SCF-flox mice were generated and transplanted with human donor CD34^+^ cells (**Figure 6D**). These recipient mice constitutively lack SCF expressed by murine *Lepr^+^* MSCs. Engraftment kinetics are severely blunted in B6.RgW41CD47;*Lepr-cre^ki/+^;SCF^fl/fl^*mice compared to controls (**Figure 6E**), suggesting impaired engraftment of donor leukocytes in the absence of *Lepr^+^* MSC-derived SCF. Of note, MSC numbers in B6.*RgW41CD47;Lepr-cre^ki/+^;SCF^fl/fl^* mice remain unaltered compared to controls (**Figure 6F**) further strengthening the evidence that lack of the molecular signal, but not absence of the cell type mediates impaired engraftment of human donor HSPCs. Blunted leukocyte chimerism in the blood is mirrored by reduced human leukocyte engraftment in the BM based on lower numbers of human lymphocytes but also cells of the myeloid lineage (**Figure 6G**). Finally, human HSPCs are almost non-detectable in the BM (**Figure 6 H**) and spleen of B6.*RgW41CD47;Lepr-cre^ki/+^;SCF^fl/fl^*mice (**Figure 6I**), suggesting persistence of only few human progenitors in the absence of considerable human HSC engraftment. Together, this shows that murine *Lepr^+^*MSC-derived SCF is required for human LT-HSC engraftment in xenotransplantation into KIT-deficient mice, suggesting not only cross-species communication but a highly conserved mechanism of the SCF-KIT axis for HSC maintenance across species.

## DISCUSSION

A major obstacle to understanding how the microenvironment regulates human HSCs *in vivo* has been the lack of experimental systems that combine robust humanisation with genetic tractability of the host niche. KIT receptor-mutant recipient mice show much improved engraftment of human immune cells after HSPC xenotransplantation compared to parental strains (Cosgun et al., 2014; Rahmig et al., 2016; Mende et al., 2019; Coppin et al., 2021). Here we provide evidence that long-term engraftment of human HSCs is also much improved in NSGW41 mice. We show a 100-fold improved engraftment for human donor leukocytes compared to irradiated NSG recipient mice and efficient engraftment in secondary recipients. The relative contribution of human donor HSCs to the stem cell pool is much increased in NSGW41 mice compared to irradiated NSG mice and endogenous murine stem and progenitor cells are numerically decreased in NSGW41 only. This provides compelling evidence for a competitive advantage for incoming, KIT-proficient human donor cells compared to endogenous, KIT-mutant murine stem and progenitor cells.

However, the utility of NSGW41 mice for mechanistic studies of the bone marrow niche is limited by the restricted availability of compatible genetic tools. Consequently, most studies have focused on characterising human hematopoiesis itself rather than interrogating how defined niche populations regulate human stem-cell behaviour *in vivo.* To address this need, we establish B6.RgW41CD47 mice which allow for human HSC and leukocyte engraftment at a level comparable to NSGW41 mice. By placing the W41 KIT mutation and CD47 deficiency onto a C57BL/6 background, this model becomes directly compatible with the extensive repertoire of existing Cre-driver, reporter, inducible deletion and disease-model strains available in the C57BL/6 genetic ecosystem. As a result, B6.RgW41CD47 mice can now be crossed with niche-specific genetic tools to selectively visualise, manipulate or ablate candidate niche cell populations and signalling pathways *in vivo*, providing a genetically tractable platform for studying human HSC–niche interactions.

Bone marrow stromal niche cells regulate stem and progenitor function in steady state (Comazzetto et al., 2021), leukemia (Urs et al., 2024), infection (Johnson et al., 2020), and aging (Matteini et al., 2021). Thus, identifying niche cells and molecular factors that control human HSC function is of high importance for the development of new therapeutic strategies in the treatment of blood cancers or other bone marrow failures. Many functional features of human and murine blood cell regulation and differentiation are comparable between the two species including hallmarks of aging of stem cells and their niche (Matteini et al., 2021) but interactions between transplanted human HSCs and the murine bone marrow niche have not been investigated to date. Based on inherent difficulties, it remains a challenge to study intra-organ communication regulating human HSC function through their native microenvironment *in situ*. Therefore, surrogate models are developed, and a new and promising research field has been opened to engineer human hematopoietic niches *in vitro* and *in vivo* (Dupard et al., 2020). However, the most promising models require subcutaneous transplantation into mice, omitting the advantage of reducing animal experiments and still relying on cross-species reactivity of cytokines and growth factors. Moreover, mainly MSCs and ECs are currently integrated in these models, reducing niche complexity significantly (Sommerkamp et al., 2021). Thus, xenotransplantation remains a viable option to learn about physiological relevance of cellular and molecular factors regulating human HSC function *in vivo*.

The microenvironment responds to incoming human donor cells with the expansion of MSCs. Molecular analysis revealed that phenotypic CD51^+^ Pdgfra^+^ MSCs comprise adipo- and osteo-primed MSCs (Dolgalev and Tikhonova, 2021; Matsushita et al., 2022) before humanization. However, after transplantation, the molecular signature indicates an almost entire shift to adipo-primed *Lepr^+^* MSCs that express hematopoiesis-supporting cytokines, indicating their critical role for HSC function. Consistently, osteogenic differentiation potential is significantly decreased. This finding translates into a bone loss phenotype comparable to what is seen in human patients after bone marrow transplantation (Schulte and Beelen, 2004). Consistent with our results, this feature has been associated with impaired osteoblast differentiation in human bones after transplantation (Lee et al., 2002). We conclude that while MSCs are a continuous population, there is a transcriptional and functional segregation into adipo- and osteo-primed MSCs. After xenotransplantation, the ratio of adipo- to osteo-primed MSCs increases and adipo-primed MSCs show transcriptional alterations that suggest an even stronger adipogenic transcriptional profile. Such plastic adaptation of MSC identity has also been suggested to occur in the context of injury (Zhou et al., 2017; Matsushita et al., 2020), and may provide the necessary flexibility to stromal niche components to balance the needs between stem cell support versus bone homeostasis.

Niche cells provide factors crucial for HSC maintenance and continuous function. *Lepr^+^* MSCs regulate endogenous murine HSC function via SCF (Ding et al., 2012) but also express further important factors such as CXCL12, Angpt1 or pleiotrophin (Pinho et al., 2013; Nakatani et al., 2023). MSCs, adipocytes, and endothelial cells are the main sources of SCF, of which *Lepr^+^* MSCs express by far the highest levels of *Scf* transcripts in the BM (Ding et al., 2012; Mende et al., 2019; Fröbel et al., 2021). Endothelial cell-derived SCF expression is not confined to the bone marrow and regulates serum protein levels but not bone marrow HSCs (Matsuoka et al., 2023) and ECs but not *Lepr^+^* cells in the spleen are crucial for SCF production for the induction of extracellular hematopoiesis (Inra et al., 2015). Thus, MSCs are a key source of regulatory SCF in the bone marrow and constitute an important niche component involved in the regulation of murine HSCs and progenitors. Importantly, an MSC population expressing CD51 and PDGFRɑ has been identified in human fetal bone marrow which is enriched for niche factors and supports human HSC expansion *in vitro*, likely representing the human equivalent of murine *Lepr^+^* MSCs (Pinho et al., 2013), suggesting conservation of the niche and its molecular factors important for HSC function between mice and humans.

Using our novel B6.RgW41CD47 mouse model in combination with a LepR-Cre deleter strain we show that deletion of LepR+ MSCs leads to mobilization of human HSCs from the bone marrow to the spleen, evidencing the critical role of LepR+ MSCs for the maintenance of human HSCs specifically in the mouse BM niche. Furthermore, the celltype specific knockout of SCF from *Lepr^+^* MSCs in B6.RgW41CD47 led to reduced engraftment and maintenance of human HSCs in the long-term, identifying the cell type but also the molecular signal that accounts for human HSC regulation. Thus, we demonstrate that our novel *in vivo* platform allows the identification of a fundamental regulatory process that is functional across species-barriers.

Overall, our data provide evidence for functional cross-species niche - HSC interactions in a mouse model with an unperturbed bone marrow microenvironment. The novelty is three-fold: We have shown proof of principle that human HSC engraftment and function is efficiently regulated by the murine niche, and, inversely, that the murine niche undergoes remodelling in response to incoming human HSCs on a cellular and molecular level, and third, we established the B6.RgW41CD47 strain as a versatile and genetically tractable humanised mouse platform that bridges the gap between conventional xenotransplantation models and sophisticated mouse genetics. By enabling direct manipulation of the host microenvironment while maintaining robust human hematopoietic engraftment, this system represents an *in vivo* platform to evaluate the functional significance of specific niche-derived factors for the modulation of human HSC function within a physiological niche context. Leveraging this approach may facilitate the identification of niche-targeted interventions to improve human HSC function and transplantation outcomes.

## MATERIAL AND METHODS

### Mice

Mouse generation: **NSG** mice were purchased from Jax #005557. **NSGW41** mice were generated in house (Cosgun et al., 2014; Rahmig et al., 2016). **B6.RgW41CD47** (Rag-2^ko/ko^;Il2rg^ko/ko^;Kit^W41/W41^CD47^ko/ko^) mice were generated by breeding RgW41 (Rag-2^ko/ko^;Il2rg^ko/ko^;Kit^W41/W41^, (Perçin et al., 2025)) with B6.129S7-CD47tm1Fpl/J ((Lindberg et al., 1996), Jax #003173). **B6.RgW41CD47;Lepr^cre/wt^;R.DTR^ki/ki^** were generated by breeding B6.RgW41CD47 mice with Lepr^cre/wt^ mice (Lepr^cre^, B6.129-Leprtm2(cre)Rck/J (DeFalco et al., 2001) Jax #008320). Subsequently B6.RgW41CD47;Lepr^cre/wt^ males were bred with R.DTR^ki/ki^ female mice (C57BL/6-Gt(ROSA)26Sortm1(HBEGF)Awai/J (Buch et al., 2005), Jax #007900). **B6.RgW41CD47;Lepr^cre/wt^;SCF^fl/fl^** (SCF^fl^ (Ding et al., 2012), Jax #017861) mice were generated using an analogous breeding strategy. All genotyping details are provided in table S1.

### Animal procedures

All mice were bred and kept under specific pathogen-free conditions in separated ventilated cages in the animal facility of the TU Dresden or Leibniz Institute on Aging in Jena providing a 12 h/12 h light/dark cycle (7am-7pm) at a temperature of 22 ± 2dC and 55 ± 10% humidity (40–75% tolerance limit). All animal experiments (Mus musculus) were performed in accordance with German animal welfare legislation and were approved by the relevant authorities: Landesdirektion Dresden or the Thüringer Landesamt für Verbraucherschutz (TLV). For **transplantation** CD34^+^ human HSPCs were isolated using dual magnetic beads enrichment according to the manufacturer’s instructions (Miltenyi Biotech) <u>(Cosgun et al.,</u> <u>2014)</u>. Purities >95% were considered acceptable. Contaminating T cell frequencies were routinely below 1%. 5×10^4^ HSPCs were transplanted intravenously into non-irradiated NSGW41 or irradiated NSG (2Gy) recipients. Mice between 5 and 17 weeks of age were transplanted with 5×10^4^ or the indicated number of CD34-enriched human cord blood cells via retroorbital injection. NSG mice received an irradiation dose of 2 Gy prior to injection (X-Ray source, MaxiShot, Yxlon; gamma cell 40 exactor, Best Theratronics). All other mice were not conditioned before transplantation. **ELDA** Limiting dilution analysis was performed by transplanting 1×10^3^, 1×10^4^ and 5×10^4^ CD34-enriched cord blood cells into non-conditioned NSGW41 or irradiated NSG mice. Mice with a human CD45 chimerism >5% were considered repopulated. Frequency of scid-repopulating cells was calculated using the ELDA software (https://bioinf.wehi.edu.au/software/elda/). For determination of functional stem cell frequency within primary recipients, secondary transplantations of 1×10^6^, 2×10^6^ or 4×10^6^ sorter-purified human CD45^+^ cells from bone marrow of CD34^+^ CB-engrafted NSGW41 or irradiated NSG mice into the same strains were performed. After 30 weeks bone marrow chimerism was analyzed and chimerism >2% was considered positive for human engraftment. The number of functional HSCs was calculated by dividing the total number of human CD45^+^ cells per 2 femora in primary recipients by the frequency of scid-repopulating cells calculated by ELDA of secondary recipients. **Diphtheria toxin** was applied 21 weeks after human CD34^+^ CB injection. Mice were injected for two consecutive days intravenously with 100 ng of unnicked Diphtheria Toxin from *Corynebacterium diphtheria* (List Biological Laboratories) diluted in sterile PBS. On day 4 after niche cell depletion mice were analyzed.

### Human material, Ethics

Cord blood samples were obtained from the DKMS Stem Cell Bank, Dresden, and from the Department of Obstetrics, Jena University Hospital, and were used in accordance with the guidelines approved by the ethics committees of Dresden University of Technology, and University Clinics Jena. After density centrifugation (BioColl 1.077 g/ml, Bio&Sell) cord blood samples were magnetically enriched for CD34^+^ cells according to manufacturer’s instructions (Miltenyi Biotec).

### Three-dimensional micro–computed tomography

After sacrifice, humeri were collected and fixed in 4% paraformaldehyde in PBS for 48 h. The bones were then briefly washed and stored in 50% ethanol until scanning. Changes in bone microarchitecture were analyzed across the entire humeri using a vivaCT 40 system (Scanco Medical, Brüttisellen, Switzerland) at an X-ray energy of 70 kVp (114 μA, 200 ms integration time) and an isotropic voxel size of 10.5 μm. The system was routinely calibrated using hydroxyapatite phantoms for density and geometric accuracy. Predefined Scanco scripts were used for evaluation. Customized data analysis: The segmentation of bone from background was performed by using texture based segmentation using filters following (Malik et al., 2001) as this method has previously been demonstrated to be highly accurate at separating bone from background in microCT images (Svensson et al., 2017; Kresinsky et al., 2019). The average bone density was calculated by computing the average intensity of the segmented bone directly from microCT data. For visualization of the bone density we used Napari’s 3D viewer (https://napari.org/). To avoid that extreme values skewing the color scale, we find a max value that corresponds to the 99.99^th^ percentile of all the values across the bone intensities. Any voxels that have intensities higher than this threshold are then capped at the 99.99^th^ percentile and are used as the max for the color map. This ensures that the color scales between separate bones are comparable.

### Flow cytometry

Hematopoietic and niche cells were stained for 40 min on ice as described before (Cosgun et al., 2014; Mende et al., 2019). Briefly, cell suspensions were stained in PBS/5% FCS with addition of blocking agents (rat IgG, mouse IgG (Dianova), CD16/32 (eBioscience)) and the indicated antibodies. A full list of antibody panels is provided in **Table S2.** Samples were acquired or sorted on LSRII, Fortessa or Aria II cytometers (BD Biosciences). All data was analysed using FlowJo software (BD Biosciences).

Mean MFI values were derived from the statistics function of FlowJo. Fold change of MFI was calculated individually for each experiment with the MFI of control NSG or B6/SJL mice serving as reference values.

### Cell suspensions

For analysis of hematopoietic cells from **bone marrow** one femur was crushed using a mortar and pestle. **Stromal cell populations** were prepared from the second femur as previously described (Mende et al., Blood, 2019). Bone marrow and bone fractions were collected and digested separately. Femurs were flushed with 4ml PBS/5% FCS using a 23G needle and briefly crushed in 2ml PBS/5% FCS and cut into small pieces. The retained cell suspension was combined with the flushed cell fraction to obtain the central BM = ’marrow fraction’. Compact bone pieces were collected in a 2 ml centrifuge tube and further processed to receive the ‘endosteal cell fraction’. After red blood cell lysis (ACK lysing buffer, Gibco) the marrow fraction was incubated with 1ml Collagenase Type I solution (3mg/ml Collagenase Type I (Worthington) in DMEM with 10% FCS and 500 μg/ml DNAse I (Sigma)) for 40 min at 37 °C at continuous mild shaking (20 rpm). In parallel, bone pieces were incubated in 1 ml Collagenase Type I solution (see above) for 20 min at 37 °C with harsh shaking at 1350 rpm. After digestion was stopped with 10 ml PBS/5% FCS, the suspensions were filtered through 100 μm mesh to remove cell clumps and bone pieces and processed further.

### SCF protein measurement

SCF concentrations in serum and bone marrow supernatant were measured using the Quantikine mSCF ELISA kit (R&D) according to the manufacturer’s instructions. Bone marrow supernatant was prepared by crushing one femur in 300ul PBS, the cell suspension was spun down for 5min at 400g and supernatant collected. Serum was obtained by centrifuging blood samples for 15min at 7600g in the presence of a few coagulation activator beads from the S-Monovette (Sarstedt) and then taking the supernatant.

### Cell culture

For CFU-F assays 1500-2000 CD45^-^CD51^+^PDGFR^+^Sca-1^-^ bone marrow MSCs were sorter purified and seeded onto gelatine-coated 6-well plates in 1 ml MSC medium (aMEM, 10% FCS, 0.1% b-mercaptoethanol, 1% Pen/Strep) and 250 ul of pre-conditioned MSC medium. After 16 to 20 days of culture, colonies were stained with Giemsa (1:20) and counted using a conventional light microscope.

### Adipogenic differentiation

was conducted slightly modified as in (Percin et al. 2025). After digestion 20% of the ‘marrow fraction’ cells of 1 femur were plated in 200 μl MSC medium for 3 days, then medium was renewed and adipogenic induction factors added (100 nM dexamethasone (D2915-100mg, Sigma Aldrich), 10 ug/ml insulin (Sigma Aldrich I5500-100 mg), 125 uM indomethacin (Sigma Aldrich I7378), 0.5 mM IBMX (Sigma Aldrich I5879) and 1 nM triiodothyronine (Sigma Aldrich T2877). After 3 more days, medium was renewed again only adding 100 nM dexamethasone and 10 ug/ml insulin to maintain the adipogenic differentiation. After 4 more days, cells were washed with PBS and fixed with 4% PFA for 30 min, washed with PBS and equilibrated in 200 μl 60% isopropanol. For staining of lipid droplets in the cells 200 ul of freshly mixed 2 parts of 0.22 μm filtered Oil red O solution (Stock: 0.3% Oil red O powder in 100% isopropanol, Sigma Aldrich O0625-25G) with 1 part water was added to the wells for 30 min at room temperature. Cells were washed with 60% isopropanol and images taken by BZ-9000E (Keyence). Next, 200 μl of 100% isopropanol was used to destain for 30 min and subsequently, 100 ul destaining solution per sample was transferred to 96-well F-bottom plate for absorbance measurement at 510 nm wavelength (TECAN Infinite M1000 Pro).

### Osteogenic differentiation

was conducted slightly modified as in (Percin et al. 2025). Briefly, after digestion 20% of the ‘marrow fraction’ cells of 1 femur were plated in 200 μl osteogenic differentiation medium containing ascorbic acid-free a-MEM (Gibco, A1049001), 10% FCS, 1% Pen/Strep, 10 mM β-glycerophosphate disodium salt hydrate (Sigma-Aldrich, G9422-10G) and 50 μg/ml 2-phospho-L-ascorbic acid trisodium salt (Sigma 49752-10G) in 96 well plates (Cell Star). Medium was replaced twice weekly. On day 14, cells were washed with PBS and fixed with 4% PFA for 30 min, washed with PBS, and stained with 200 μl Alizarin Red Solution (Merck #2003999) for 30 min at room temperature. Wells were washed with PBS and images were taken by BZ-9000E (Keyence). Afterwards, wells were incubated with 200 μl destaining solution (70% water, 10% acetic acid, and 20% methanol) for 30 min at room temperature and subsequently, 100 ul destaining solution per sample was transferred to 96-well F-bottom plate for absorbance measurement at 450 nm wavelength (TECAN Infinite M1000 Pro). For SCF cultures 15.000 human CD34^+^ CB cells were plated in 96-well U-bottom plates (Nunc) in 200 ul StemSpan medium (StemCell Technologies) containing 10 ng/ml human TPO (PeproTech), 50 ng/ml human FLT3L (PeproTech), 500 ng/ml Angpt-like protein 5 (ANGPTL, Miltenyi Biotec) and either 20 or 100 ng/ml of murine or human SCF (PeproTech). Medium was changed and cells transferred to 24-well plates, 12-well plates and 6-well plates every 3 days and cell expansion measured after 10 days in total.

### Statistical analysis

Data were analyzed using Welch’s t-test, which maintains robustness against unequal variances and unbalanced sample sizes. Other statistical tests are indicated in the figures legends.

### Molecular analysis and NGS library preparation

For **bulk sequencing** Ter119^-^Lin^-^CD45^-^CD31^-^CD51^+^PDGFR^+^Sca-1^-^ bone marrow MSCs were sorter purified from humanized, irradiated NSG (18 wks) or NSGW41 (23 wks) mice (5500 - 7500 cells per mouse) and non-transplanted controls (500 - 1700 cells, 2 NSGW41 mice pooled per sample). RNA was prepared using the miRNeasy Micro Kit (Qiagen). Total RNA was eluted in 10ul and 5ul were used for library preparation for next generation sequencing. cDNA was synthesized from 5 µl total RNA using the SmartScribe reverse transcriptase (Takara Bio, SMARTer HV Kit) with a universally tailed poly-dT primer and a template switching oligo followed by amplification for 12 cycles with the Advantage 2 DNA Polymerase (Takara Bio). After ultrasonic shearing (Covaris LE220), amplified cDNA samples were subjected to standard Illumina fragment library preparation using the NEBnext Ultra DNA library preparation chemistry (New England Biolabs). In brief, cDNA fragments were end-repaired, A-tailed and ligated to indexed Illumina Truseq adapters. The resulting libraries were PCR-amplified for 15 cycles using universal primers, purified using XP beads (Beckman Coulter) and then quantified with the Fragment Analyzer. Final libraries were equimolarly pooled and subjected to 75-bp-single-end sequencing on the Illumina Nextseq 500 platform, resulting in on average 35 mio. fragments.

For **single cell RNAseq** single CD45^-^CD51^+^PDGFR^+^Sca-1^-^ bone marrow MSCs were isolated from humanized NSGW41 24 or 25 weeks after humanization and processed based on a published protocol (Picelli et al., 2013). Single cells are sorted with FACS into a 96 well plate containing 2 µl of nuclease free water with 0.2% Triton-X 100 and 4 U murine RNase Inhibitor (NEB), spun down and frozen at -80°C. After thawing the samples, 2 µl of a primer mix is added (containing 5 mM dNTP (Invitrogen), 0.5 µM dT-primer*, 4 U RNase Inhibitor (NEB)). RNA is denatured for 3 minutes at 72°C and the reverse transcription is performed at 42°C for 90 min after filling up to 10 µl with RT buffer mix for a final concentration of 1x superscript II buffer (Invitrogen), 1 M betaine, 5 mM DTT, 6 mM MgCl2, 1 µM TSO-primer*, 9 U RNase Inhibitor and 90 U Superscript II. After synthesis, the reverse transcriptase is inactivated at 70°C for 15 min. The cDNA is amplified using Kapa HiFi HotStart Readymix (Peqlab/Roche) at a final 1x concentration and 0.1 µM UP-primer* under following cycling conditions: initial denaturation at 98°C for 3 min, 22 cycles [98°C 20 sec, 67°C 15 sec, 72°C 6 min] and final elongation at 72°C for 5 min. The amplified cDNA is purified using 1x volume of hydrophobic Sera-Mag SpeedBeads (GE Healthcare) rebuffered in a buffer consisting of 10 mM Tris, 20 mM EDTA, 18.5 % (w/v) PEG 8000 and 2 M sodium chloride solution. The cDNA is eluted in 12 µl nuclease free water and its concentration is measured with a Tecan plate reader Infinite 200 pro in 384 well black flat bottom low volume plates (Corning) using AccuBlue Broad range chemistry (Biotium).

For library preparation up to 700 pg cDNA in 2 µl is mixed with 0.5 µl Tagment DNA Enzyme and 2.5 µl Tagment DNA Buffer (Nextera, Illumina) and tagmented at 55°C for 5 min. Subsequently, Illumina indices are added during PCR (72°C 3 min, 98°C 30 sec, 12 cycles [98°C 10 sec, 63°C 20 sec, 72°C 1 min], 72°C 5 min) with 1x concentrated KAPA Hifi HotStart Ready Mix and 0.7 µM dual indexing primers. After PCR, libraries are quantified with AccuBlue Broad range chemistry, equimolarly pooled and purified twice with 1x volume of rebuffered Sera-Mag SpeedBeads. The library pools were sequenced with 75-bp-single-end on the Illumina Nextseq500 platform, aiming at an average sequencing depth of 0.5 mio. fragments per cell. dT-primer: C6-aminolinker-AAGCAGTGGTATCAACGCAGAGTCGACTTTTTTTTTTTTTTTTTTTTTTTTTTTTTTVN, where N represents a random base and V any base beside thymidine; TSO-primer: AAGCAGTGGTATCAACGCAGAGTACATrGrGrG, where rG stands for ribo-guanosine; UP-primer: AAGCAGTGGTATCAACGCAGAGT

### Bioinformatic analysis

#### Availability of code and data

Raw sequencing files and feature-by-sample (bulk) or feature-by-cell (single-cell) count matrices for both single-cell and bulk RNA-seq data have been deposited in the Gene Expression Omnibus under accession numbers GSE322658 (token: etqrouckxvkfzud) and GSE322746 (token: enujemcaftqrtsf). During revision, processed data objects generated during the computational analysis workflow (e.g., Seurat objects containing single-cell data) are available at https://tsbnc.dkfz.de/index.php/s/cn6HjX4NAPqkNr8 (password: MSC&data&2026!). Upon publication, raw and processed data will be made publicly available. Scripts used for bulk RNA-seq analyses and downstream single-cell RNA-seq analyses are available at https://github.com/jome1994/Niche_MS. Scripts used for preprocessing and core single-cell RNA-seq analysis steps are available at https://github.com/Iwo-K/NSGW41-HSC-niche-scRNASeq.

Code for the analysis of the bone density is available at https://github.com/applied-systems-biology/Froebel_CT_data_analysis and the data used for analysis at https://doi.org/10.5281/zenodo.18849011.

#### Analysis of bulk RNA-seq data

Single-end bulk RNA-sequencing reads (stored in FASTQ format) were aligned to the mouse reference genome (mm39) using STAR (v2.7.10a;(Dobin et al., 2013)). Only uniquely mapping reads were retained. Gene-level counts were generated from the resulting BAM files using featureCounts (Subread v1.6.5,(Liao et al., 2014)). Reads were assigned to exonic regions based on the Mus musculus GRCm39.108 gene annotation (GTF), summarized at the gene_id level. Counting was performed jointly across all samples to produce a single gene-by-sample count matrix, in which rows correspond to genes and columns to individual samples. Versions of R packages used for the workflow described in the following paragraphs of this section can be extracted from the renv.lock file provided with the code. The list includes packages from the Bioconductor ecosystem (Huber et al., 2015). The gene-level count table was imported into R. The table was separated into a gene annotation component (first five columns) and a raw count matrix (remaining columns). Genes with very low detection were filtered out by removing features with non-zero counts in fewer than three samples. Ensembl gene IDs were translated to gene symbols using AnnotationHub resources (Ensembl mm39, version 105). Where necessary, duplicate symbols were resolved by generating unique feature names. Entrez IDs were additionally retrieved based on the mapped gene symbols using the org.Mm.eg.db database. The filtered count matrix was then relabeled to use unique gene symbols as row names. For exploratory data analysis and visualization, counts were normalized using the variance stabilizing transformation (VST) implemented in DESeq2 (Love et al., 2014). Principal component analysis (PCA) was performed on the transformed data using the 500 most variable genes.

Genotype-specific differential expression analyses were performed independently to quantify and compare the number of significantly up- and downregulated genes upon humanization (Table S3). For each genotype, a separate DESeqDataSet was constructed. Prior to model fitting, genes were filtered. A gene was kept if it had counts ≥10 in ≥(n−1) samples within at least one condition, where n is the number of replicates per condition. Conditions were then compared independently within each genotype using DESeq, followed by Wald test and filtering of test results using a significance threshold of adjusted p < 0.05.

For gene set enrichment analysis (GSEA), shrinkage-estimated log2 fold changes were derived from a single joint model containing all samples from both genotypes. A unified DESeqDataSet was fitted across conditions with dispersion estimation performed once for the full dataset (DESeq, with outlier replacement disabled via minReplicatesForReplace=Inf). Log2 fold changes were then extracted separately for each genotype-specific treatment contrast (NSG_Tx vs NSG_no_Tx and NSGW41_Tx vs NSGW41_no_Tx) using lfcShrink with adaptive shrinkage (“ashr”). This approach ensured that (i) fold changes were computed for the same gene set in both genotypes without genotype-specific prefiltering, and (ii) shrinkage estimation was based on a common dispersion framework, resulting in directly comparable ranked gene vectors for downstream enrichment analyses.

#### Gene set enrichment analysis for bulk and single-cell RNA-seq data

Versions of R packages used for the workflow described in this section can be extracted from the renv.lock file provided with the code.

To identify biological processes associated with transcriptional changes, we performed Gene Set Enrichment Analysis (GSEA) using shrinkage-adjusted log₂ fold. Rather than testing genes individually, GSEA evaluates whether predefined groups of genes - referred to as *gene sets* - show coordinated up- or downregulation. In this study, gene sets were defined using the Gene Ontology (GO) Biological Process collection (Gene Ontology Consortium, 2021). Each gene set corresponds to a GO term, which represents a curated functional annotation describing a biological process (e.g., “regulation of phosphorylation”). Thus, when referring to “significant GO terms,” we mean the functional annotations of gene sets that were found to be significantly enriched.

Gene set enrichment analysis was conducted using the clusterProfiler framework (Wu et al., 2021) together with GO annotations from org.Mm.eg.db. Genes were ranked by their shrinkage-adjusted log₂ fold change without additional prefiltering. To avoid artificial ranking bias caused by identical fold-change values, tied genes were randomly ordered. Gene symbols were mapped to Entrez identifiers, and enrichment was computed using gene sets containing between 10 and 800 genes. Adjustment of p-values was done using the Benjamini-Hochberg method.

For each enriched gene set, GSEA identifies a subset of genes known as the *leading edge genes*. These are the genes within the gene set that contribute most strongly to the enrichment score—that is, the subset of genes driving the coordinated up- or downregulation signal. Biologically, the leading edge can be interpreted as the core group of genes responsible for the enrichment of a given functional annotation.

To focus on robust biological signals and reduce redundancy among overlapping GO terms, only gene sets with an adjusted p-value < 0.05 were retained. Because many GO gene sets partially overlap in gene composition and may describe closely related biological processes, we quantified similarity between enriched gene sets based on their leading edge genes. For each pair of gene sets, we calculated the Jaccard index (intersection divided by union of leading edge genes), which measures the fraction of shared genes between two sets.

We then constructed a similarity network in which each node represents a significantly enriched gene set (i.e., its associated GO term), and edges connect pairs of gene sets whose leading edge overlap reached or exceeded a Jaccard index of 0.5. This threshold ensured that only gene sets sharing at least half of their leading edge genes were considered strongly related. The resulting network consisted of well-separated subnetworks representing groups of highly overlapping and functionally related biological processes.

Using this network structure, gene sets were iteratively clustered into consensus “umbrella sets.” Operationally, clustering was performed by identifying connected components within the similarity network. Starting from individual enriched gene sets, any two sets connected by an edge (Jaccard ≥ 0.5) were assigned to the same cluster. If a third gene set overlapped sufficiently with any member of an existing cluster, it was added to that cluster. This process continued until no additional gene sets met the similarity criterion, resulting in stable groups of highly overlapping functional annotations. Each umbrella set therefore represents a higher-order functional module composed of multiple related GO terms that share a substantial core of leading edge genes. Umbrella sets were manually annotated according to their dominant biological theme. Only clusters containing at least 10 unique leading edge genes in total were retained for further analysis.

For each umbrella set, the union of all its leading edge genes was used to represent the core transcriptional program underlying that functional module. To visualize the gene-level drivers of enrichment, heatmaps were generated using the combined leading edge genes across all umbrella sets. Genes were retained if their maximum absolute log₂ fold change was ≥ 0.5; if more than 70 genes fulfilled this criterion, the top 70 ranked by absolute fold change were selected. Fold-change values were displayed only for gene–umbrella set combinations in which a gene contributed to the leading edge of that umbrella set, while all other cells were left blank to emphasize module specificity. To prevent extreme values from dominating the color scale, fold changes were clipped to the 5th–95th percentile range. Genes were ordered by fold change to facilitate interpretation.

For bulk RNA-seq data, Gene Set Enrichment Analysis (GSEA) was performed separately for each genotype to identify significantly deregulated functional modules (umbrella sets) in response to humanization (Table S4). For each genotype, functional modules were filtered using a minimum of 30 leading-edge genes per module. To enable a direct comparison of transcriptional responses between genotypes, we constructed the union of all retained genes sets that were identified in either genotype. Gene sets included in this union were re-grouped into consensus umbrella sets in the following way: We connected gene sets that shared at least 50% (Jaccard-index >= 0.5) of the genes that contributed most to their enrichment scores (leading-edge subset). Violation of this criterion in at least one mouse line with both sets being significantly enriched, prevented their connection. For each genotype, we then quantified and visualized the expression changes associated with this combined set of umbrella modules. This approach ensured that functional modules uniquely enriched in one genotype were still evaluated in the other genotype, thereby allowing symmetric and unbiased comparison of pathway-level regulation across genotypes.

### Single-cell RNA-seq data analysis

Versions of software used in this paragraph can be extracted from the singularity image provided with the code.

Single-end Illumina-reads generated from the Smart-seq libraries were mapped to reference genome mm10 using STAR. Count matrices were generated using the featureCounts function from the SubRead software.

Data quality per cell was assessed in the following way: For each cell, the total number of mapped reads, fraction of reads mapped to the mouse genome, fraction mapped to ERCC spike-ins, fraction mapped to mitochondrial genes, and the number of highly expressed genes were computed. The number of highly expressed genes was defined as the count of genes with expression >50 counts per million. Cells were classified as low quality and removed if they met any of the following criteria: fewer than 100,000 mapped reads, more than 3×10⁷ mapped reads, ERCC read fraction >0.15, fewer than 900 highly expressed genes, or mitochondrial read fraction >0.2. Additionally, cells with a low fraction of reads mapping to the mouse genome were excluded. The remaining cells passing all filters were retained for downstream analysis. Library size normalisation was performed on the filtered count matrix with the scran method (Lun et al., 2016). Briefly, size factors were calculated for each cell using the scran deconvolution approach (computeSumFactors) based on counts from endogenous mouse genes. Raw counts were then divided by the corresponding cell-specific size factors to obtain normalised expression values. No pre-clustering was applied prior to size factor estimation. Dimensionality reduction and clustering were performed in Python using Scanpy (Wolf et al., 2018). Normalised counts were log-transformed (log1p) and scaled prior to principal component analysis (PCA), which was computed using 50 components. A k-nearest neighbor graph was then constructed using the first 20 principal components with 7 neighbors per cell. Community detection was carried out using the Leiden algorithm (Traag et al., 2019) at a resolution parameter of 0.45 to define transcriptionally distinct cell clusters. For visualization, a two-dimensional embedding was generated using Uniform Manifold Approximation and Projection (UMAP) based on the computed neighborhood graph. Coordinates from the UMAP embedding were used to visualize cells colored by metadata annotations and Leiden cluster identity.

Differential expression analysis was conducted in R using DESeq2 on the quality-controlled count matrix restricted to Ensembl-annotated genes. Genes with very low expression were prefiltered (mean raw count >1.5 across cells). Pairwise comparisons were performed by subsetting the data to the two conditions of interest (Humanized vs non-humanized) and fitting a negative binomial model with treatment as the design factor (∼Treat), using non-humanized as the reference level. Cell-specific size factors estimated with the scran deconvolution method were supplied for normalization. Wald tests were used to assess differential expression, and p-values were adjusted for multiple testing using the Benjamini–Hochberg method. Genes with adjusted p-values <0.1 were considered significant (Table S5). Shrunken log2 fold changes were additionally computed (lfcShrink, normal and ashr) to obtain more stable effect size estimates. For Gene Set Enrichment Analysis, all genes (not just the significant ones) were ranked based on their shrunken log2 fold changes computed by the ashr method (Table S6). Gene Set Enrichment Analysis was conducted as described in the section above.

For cell type annotation, single-cell RNA-seq profiles were mapped onto the Dolgalev & Tikhonova reference atlas (Dolgalev and Tikhonova, 2021). Count matrices of the target and reference dataset were normalized (10,000 counts per cell), log-transformed, and restricted to shared highly variable genes. The target and reference datasets were aligned in a common gene space. Target cells were projected onto the reference using a k-nearest-neighbor approach (k = 10). Cell type labels were transferred from reference to target cells by majority voting among nearest neighbors with a 60% agreement threshold. Vice versa, cluster annotations were projected from our data to the reference cells.

Upon the described processing steps and core analyses, RNA counts and cell annotations were aggregated in a Seurat object (Hao et al., 2021). Transcript counts of individual genes and expression scores of gene modules were then visualized on the UMAP-embedding using the Seurat framework.

## Supporting information

List antibodies used

Genotyping primers

DEGs of scRNAseq data

GSEA of bulk RNAseq data

GSEA of scRNAseq data

DEGs of bulk RNAseq data

## Author contributions

JF and SRa performed all experiments, analyzed data, prepared figures, and contributed to manuscript writing. JM and TH generated the gene by sample count matrix from bulk RNA-sequencing data and conducted downstream bulk analyses. They further conducted the Gene Set Enrichment Analyses for bulk and single-cell RNA-seq data and implemented a customized analytical strategy to identify functional modules of GO terms. IK and BG generated the count-matrix out of FastQ files of the SmartSeq2 experiment, performed quality controls and data normalization and UMAP-embedding, cell cluster establishment, DEG between conditions and projection of cluster-specific cell profiles onto the niche-atlas. CMS and MTF performed customized analysis of microCT data and prepared corresponding figures. SRe and AD performed library preparations and bulk and scRNA sequencing. JSH and MR performed microCT analysis and provided help with the analysis. EC performed transplantation experiments and analysis. NM contributed to the phenotyping and sorting of bulk niche cells, NH contributed to the establishment of *in vitro* functional niche cell assays and *in vivo* experiments, and GP contributed to microCT analysis. FW, AK, TG, and AP provided cord blood samples. CW conceived the study, acquired funding, designed experiments, interpreted data, and wrote the manuscript.

## Acknowledgements

The authors thank Sabrina Eichwald, Elke Meier, and Melissa Krija for excellent support in genotyping and analysis. The Core Facility and service flow cytometry as well as the animal house of the FLI are gratefully acknowledged for their technological support. We thank Sandra Hippauf for expert technical support in µCT imaging, measurements, and data analysis. This work was supported by the German Research Foundation (DFG) through TRR1278-C06, WA2837/8-1, by the Carl Zeiss Foundation (Impuls, project ID P2019-01-006), and by the DFG under Germanýs Excellence Strategy – EXC 2051 – Project-ID 390713860 (all C.W.). The FLI is a member of the Leibniz Association and is financially supported by the Federal Government of Germany and the State of Thuringia. The publication of this article was funded by the Open Access Fund of the Leibniz Association and the Leibniz Institute on Aging - Fritz Lipmann Institute (FLI), Jena, Germany. Open Access funding was enabled and organized by Project DEAL. MR and JS-H received financial support from the German Research Foundation (Deutsche Forschungsgemeinschaft, DFG) Grant Numbers DFG – SFB/TRR369 DIONE – 501752319 Project A03 and B01. JS-H acknowledges additional support from the German Federal Ministry for Research, Technology and Space (Bundesministerium für Forschung, Technologie und Raumfahrt; BMFTR) under Grant Numbers 01KC2304 and the through DFG grant number SA 4045/8-3, as well as from the Elsbeth Bonhoff Foundation (Project Number 262). BG and IK were supported by Wellcome (Z309075/Z/24/Z and Z206328/Z/17/Z). MTF received financial support from the German Federal Ministry of Research, Technology and Space within the funding program Photonics Research Germany, Project Leibniz Center for Photonics in Infection Research, Subproject LPI-BT3, contract number 13N15709.

---

**Figure S1:**
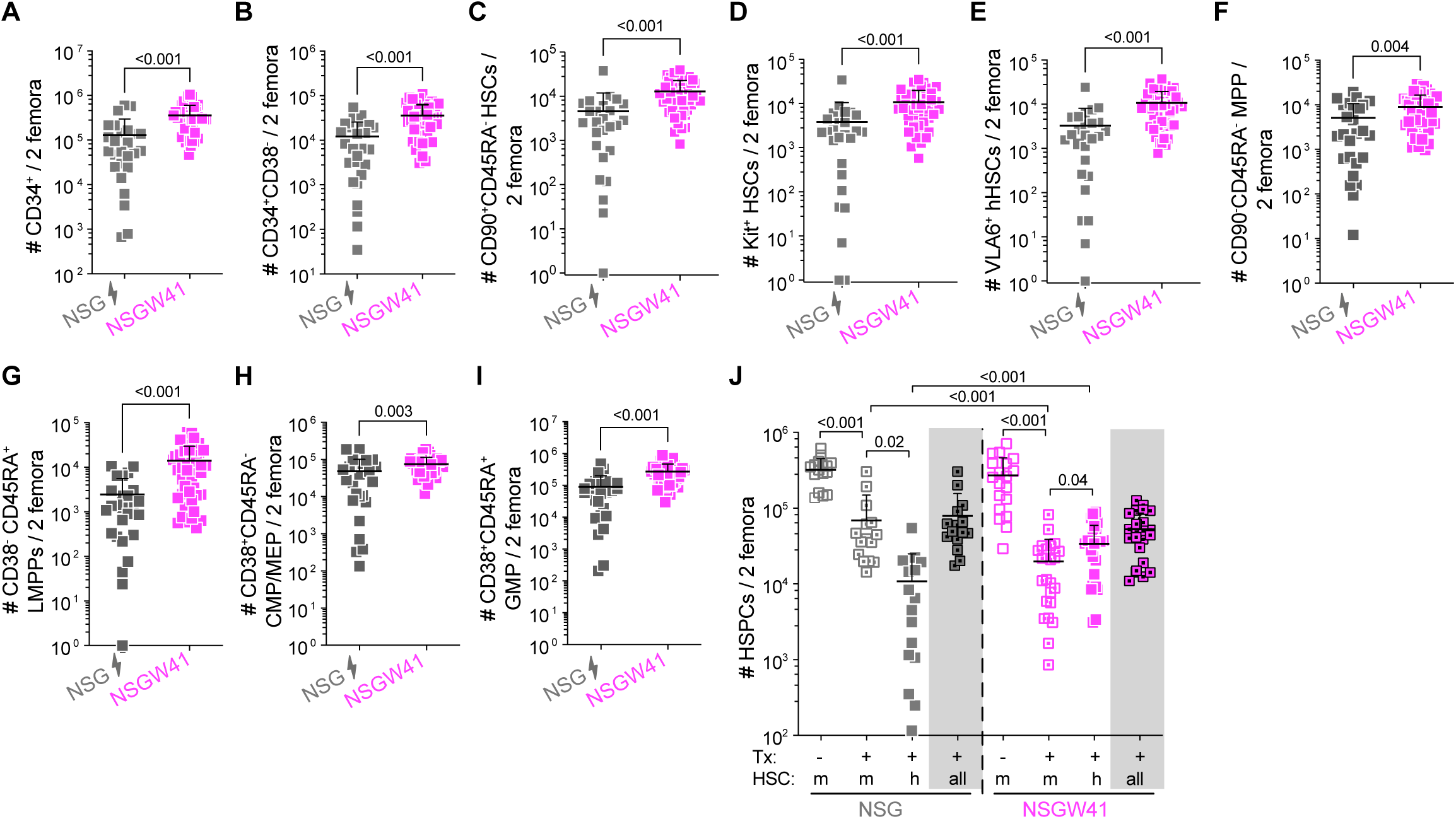
Human HSPC numbers in irrNSG and NSGW41 recipient mice. **A**-**I** Quantification of indicated human HSPC subsets gated as shown in Figure 1D according to (Notta et al., 2011; Cosgun et al., 2014; Karamitros et al., 2018; Triana et al., 2021). **J** HSPC numbers of murine (KSL) and human (CD38^-^CD34^+^) origin from the same mice as shown in Fig 1E. Statistics between different mice were calculated by Welch’s t-test, for different cells within the same mice by paired t-tests.

**Figure S2:**
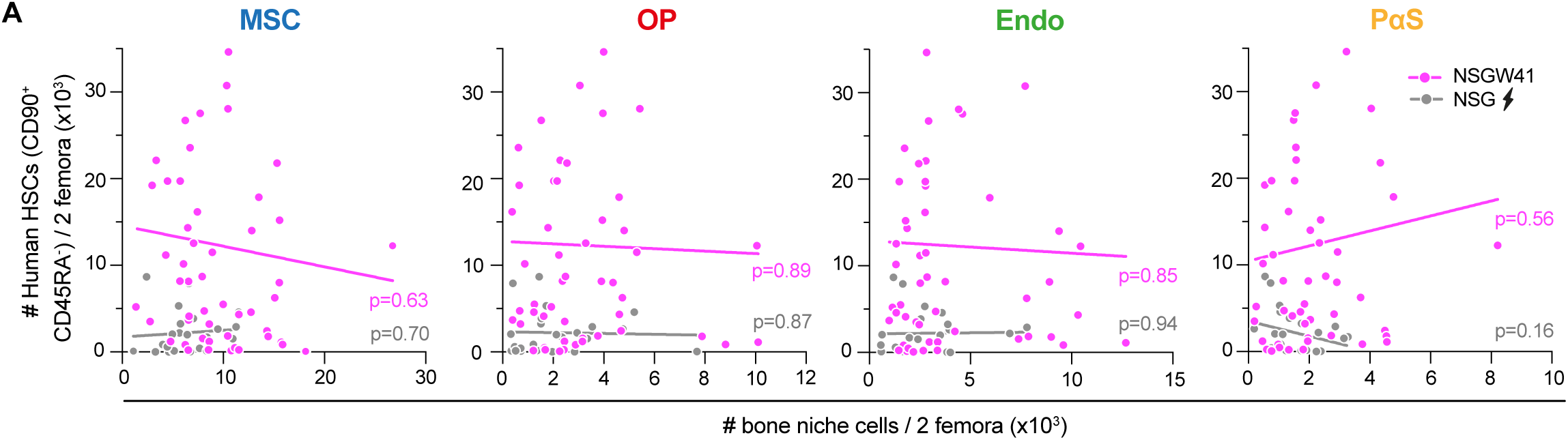
Endosteal niche space is unaltered upon humanization. **A** Correlation between the numbers of endosteal murine MSCs, osteoblasts, endothelial cells or PaS cells and engrafted human HSCs 4-35 weeks after humanization (14 independent experiments, p-values of Pearson correlation).

**Figure S3:**
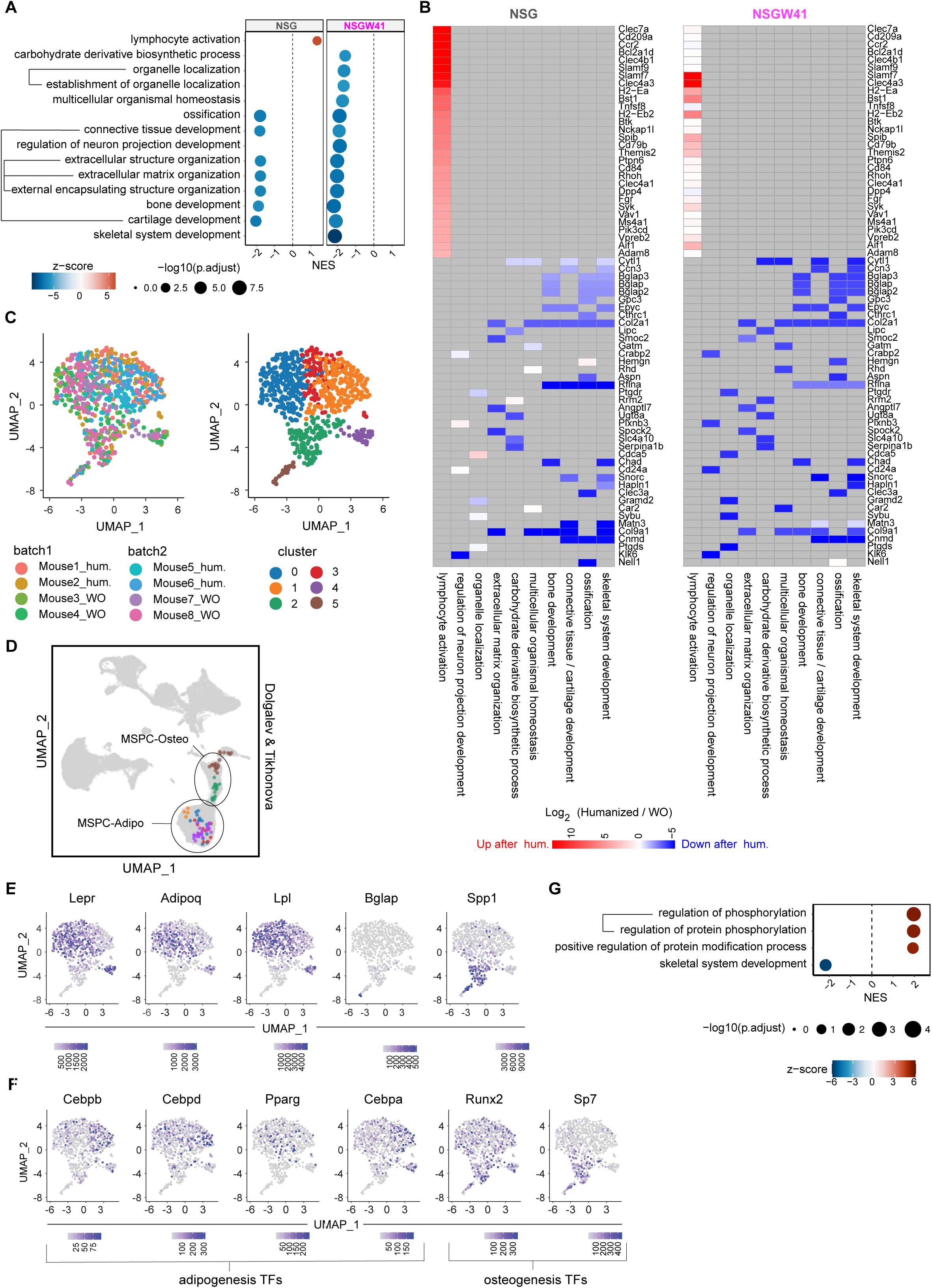
Reduced osteogenic transcriptional priming in MSCs upon humanization. **A** Dotplot of gene sets significantly deregulated upon humanization identified from bulk RNAseq data. Sets were selected by GSEA gene ontology database; category=biological processes). GSEAs were independently conducted for NSG (left) and NSGW41 (right) mice. For comparison, the same gene sets are shown for both mouse lines. Displayed sets were found to be significantly deregulated in at least one mouse line (adjusted p-value < 0.05). Dot size indicates the negative Log10 of the adjusted p-value (method: Benjamini-Hochberg). Connected gene sets share at least 50% (Jaccard-index >= 0.5) of the genes that contributed most to their enrichment scores (leading-edge subset). Violation of this overlap-criterion in at least one mouse line with both sets being significantly deregulated, prevented their connection. **B** Humanization-induced expression changes (Log2FCs) of genes per gene set module. Columns correspond to modules of gene sets identified in A. Terms that are connected in A form a module/column in B. One module corresponds to at least one gene set. Expression changes are only shown for leading edge genes (genes that strongly contribute to the enrichment scores of gene sets in A). Gray fields indicate genes that are not contributing to the enrichment of the respective gene set in any mouse line. Heatmaps include 70 leading-edge genes with largest absolute Log2FCs. For ranking, each gene was assigned to its absolute Log2FC in the mouse line in which it contributed to the significant enrichment of a gene set. In case of contributions in both mouse lines, the gene’s largest log2FC was used for ranking. 5% of smallest and largest values were clipped for better color resolution. **C** UMAP of single-cell RNA profiles colored by mouse ID (left; hum.=humanized; WO=not humanized) and by clusters identified based on the Leiden algorithm (right). **D** Niche-cell atlas from Dolgalev & Tikhonova (Dolgalev and Tikhonova, 2021). Cells in the atlas that are similar to cells of Leiden clusters in **C** are colored by the respective cluster color. **E** Normalized transcript counts of selected genes used for the identification of MSC subtypes. **F** Normalized transcript counts of transcription factors associated with adipogenic and osteogenic differentiation of MSCs as shown in Figure 3H. **G** Dotplot of gene sets significantly deregulated upon humanization in MSCs from NSGW41 mice. Sets were selected by GSEA of single-cell RNAseq data. Red color indicates upregulation in response to humanization, blue indicates downregulation.

